# Biophysical characterization reveals the similarities of liposomes produced using microfluidics and electroformation

**DOI:** 10.1101/859124

**Authors:** Michael Schaich, Diana Sobota, Hannah Sleath, Jehangir Cama, Ulrich F. Keyser

## Abstract

Giant Unilamellar Vesicles (GUVs) are a versatile tool in many branches of science, including biophysics and synthetic biology. Octanol-Assisted Liposome Assembly (OLA), a recently developed microfluidic technique enables the production and testing of GUVs within a single device under highly controlled experimental conditions. It is therefore gaining significant interest as a platform for use in drug discovery, the production of artificial cells and more generally for controlled studies of the properties of lipid membranes. In this work, we expand the capabilities of the OLA technique by forming GUVs of tunable binary lipid mixtures of DOPC, DOPG and DOPE. Using fluorescence recovery after photobleaching we investigated the lateral diffusion coefficients of lipids in OLA liposomes and found the expected values in the range of 1 μm^2^/s for the lipid systems tested. We studied the OLA derived GUVs under a range of conditions and compared the results with electroformed vesicles. Overall, we found the lateral diffusion coefficients of lipids in vesicles obtained with OLA to be quantitatively similar to those in vesicles obtained via traditional electroformation. Our results provide a quantitative biophysical validation of the quality of OLA derived GUVs, which will facilitate the wider use of this versatile platform.

## 1. Introduction

Liposomes, small aqueous compartments encapsulated by a lipid bilayer, have come a long way since their first description by Bangham and Horne in 1964 (1, 2). Today liposomes, also known as lipid vesicles, are a widely used tool in many branches of science and industry, including the pharmaceutical and cosmetics industries (3). Unilamellar liposomes of several microns in diameter, termed Giant Unilamellar Vesicles (GUVs), are especially widespread in the fields of biophysics and synthetic biology, where they are used in the bottom up construction of synthetic cells (4, 5). GUVs have also been used as model membranes for drug transport studies across lipids (6, 7) and for studying antibiotic transport facilitated by bacterial porins (8). Others have used GUVs as micro containers for chemical reactions (9).

Different methods to obtain GUV have emerged over the years which typically fall into two categories. They either generate GUVs via swelling from a solid substrate, or they are assembled from fluid interfaces (10, 11). Among the swelling approaches, a technique called electroformation found particularly widespread use (12). Many techniques of the latter category are based on microfluidics (13). A new microfluidic technique to form GUVs on chip was presented by Deshpande *et al*. in 2016 (14). Depicted in Figure 1, GUV formation occurs in a process similar to bubble blowing. At a six-way junction, an inner aqueous (IA) phase encounters a lipid-carrying 1-octanol (LO) phase. A double emulsion droplet forms and is pinched off by an outer aqueous (OA) fluid stream. Within the microfluidic chip, the double emulsion then separates, resulting in a separate GUV and an octanol droplet (14, 15). Importantly, the separation occurs automatically and the technique does not require washing or solvent extraction procedures, like other double emulsion techniques (14, 16). Since the lipids for this technique are carried by the 1-octanol phase, the method was coined Octanol-Assisted Liposome Assembly (OLA).

**Figure 1:**
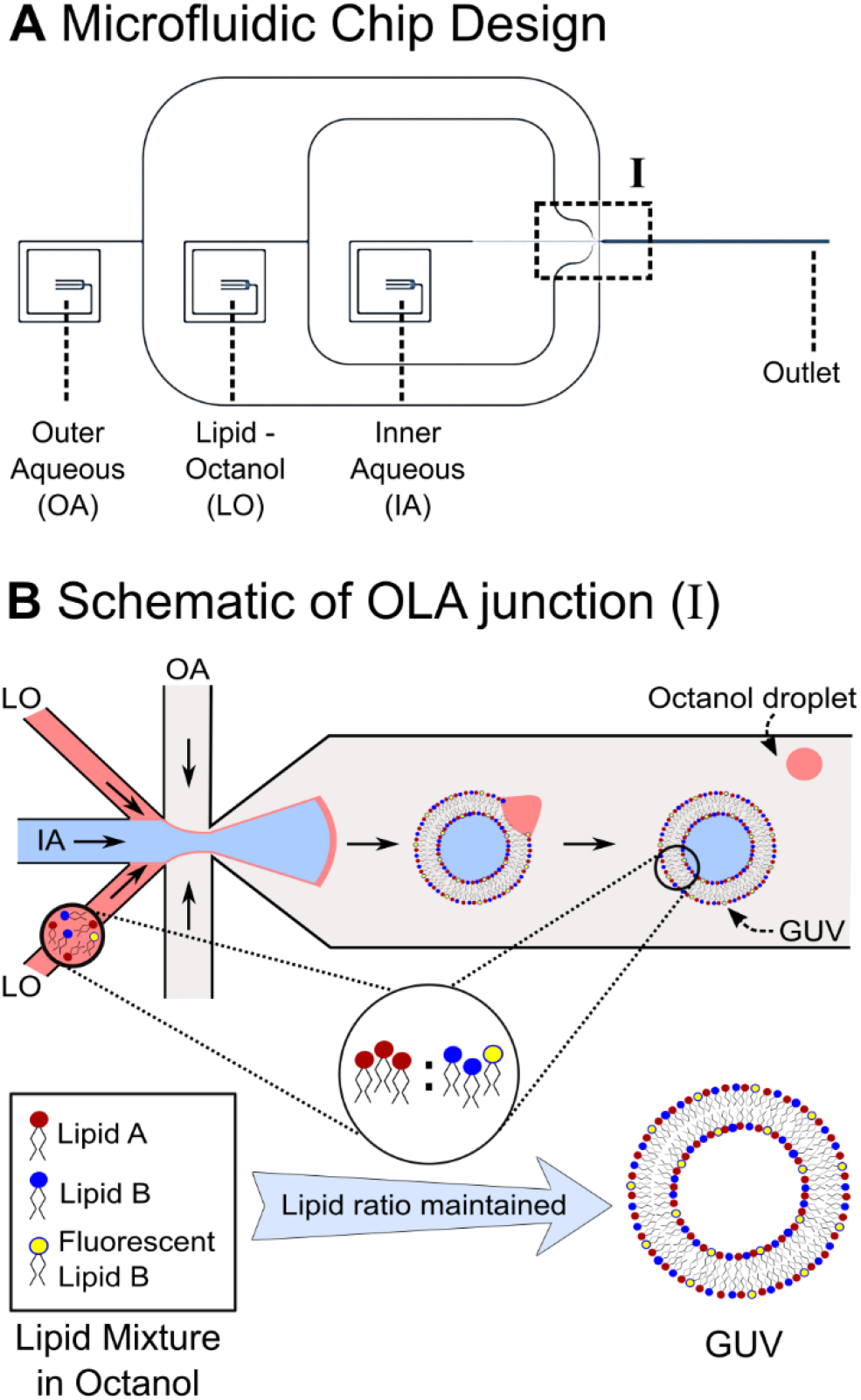
(A) Design of the microfluidic chip used to produce the liposomes. The chip has three inlets for the inner (IA) and outer aqueous (OA) and the lipid-octanol (LO) phases, respectively. The vesicles are formed at a six-way junction (I) and flow along the channel to the outlet where they can be extracted or imaged directly. (B) Schematic of the OLA junction, where liposome formation occurs. The IA fluid stream is flanked by two channels with the lipid-carrying LO phase. The OA flows pinch off double emulsion droplets. The double emulsion self-assembles downstream into a vesicle with a 1-octanol pocket attached to it, which later buds off. We propose that the lipid ratio of the lipid-octanol mixture inserted into the microfluidic chip is maintained in the GUVs produced with OLA.

Given the advantages of lab-on-chip techniques for drug development studies (17), we integrated OLA with a platform for the characterization of membrane active antimicrobials (18), as well as with a platform for the quantification of drug permeation across membranes (19). In the field of synthetic biology, mechanical division (20) and membrane tension mediated growth (21) of OLA vesicles, as well as coacervate formation within liposomes have been shown (22). Furthermore, protocols for the purification of OLA liposomes have been presented (23).

However, the capability of OLA in producing liposomes of defined lipid mixtures has not been investigated in detail previously. Theoretical and experimental studies suggest different partition coefficients of octanol into bilayers of PG, PE and PC lipids, respectively (24, 25). A lipid type with a higher affinity to octanol could therefore potentially remain in the LO phase during liposome formation. It is important to quantify that no such demixing occurs and that the membrane composition of the obtained liposome matches the lipid mixture in the LO phase. Moreover, membrane properties such as the lateral diffusion coefficients of lipids in OLA vesicles have not been compared to vesicles obtained from other GUV formation techniques. In recent years, we have gained a better understanding of the importance of membrane composition as well as lipid lateral diffusion on cellular processes. For instance, simulations by Duncan *et al*. show that clustering of Kir channel proteins is modulated by the compositional complexity as well as the lateral diffusion coefficient of the lipids in the membrane (26). Other studies have revealed that lateral lipid diffusion is rate limiting for many cellular processes (27). The proposition of different modes of lateral mobility (28), as well as the emergence of the field of lipidomics (29), furthermore highlight the increasing importance attributed to membrane composition and the lateral mobility of lipids in the membrane. When using GUVs as a tool to study proteins, precise knowledge and control of these two parameters is therefore of great importance.

In this work, we investigate the lipid composition of GUVs produced with OLA by a mean fluorescence intensity analysis. Furthermore, we measure the lateral diffusion coefficients of fluorescently labelled lipids in OLA vesicles using FRAP (Fluorescence Recovery After Photobleaching) and compare the diffusion coefficients to those obtained from vesicles generated by the established electroformation technique. To the best of our knowledge, we are the first to directly compare the lateral diffusion coefficients of GUVs obtained from a microfluidic technique to those obtained via a bulk technique. We thus provide an important biophysical characterization of liposomes produced using microfluidics, encouraging the wider uptake of these novel liposome production methods in the field.

## 2. Materials and Methods

### 2.1 Microfluidic Chip Design and Fabrication

The microfluidic chip design, depicted in Figure 1A, is a modification of the original design geometry by Deshpande *et al*. (14). The chip consists of three inlets for the inner aqueous (IA), outer aqueous (OA) and lipid-octanol (LO) phases, respectively. The LO and OA channel bifurcate and meet the IA at a six-way junction, where the liposomes are formed. As depicted in Figure 1B, the GUVs initially have an octanol pocket attached to them. The octanol typically separates from the GUV within seconds to minutes after formation, as the vesicle flows towards the outlet reservoir (14). The dimensions of our junction are scaled up by a factor of 2 compared to the original design to obtain larger liposomes than typically possible with the originally published chip design (14). The scaled-up channels furthermore lead to higher flow rates and higher liposome production rates than the original device (19). However, if operated in a high flow rate regime, the vesicles often only have approx. 30 seconds from their production until they reach the outlet, compared to up to several minutes in the original design. It should be noted, that this occasionally does not leave enough time for every single GUV to separate from its octanol pocket before it reaches the outlet. For this reason, we sometimes find GUVs with residual octanol attached in the outlet reservoir.

The microfluidic chips were fabricated from polydimethylsiloxane (PDMS) using established photo- and soft lithography techniques (19). A master mold of the structures of the microfluidic chip was produced by spin coating a thin layer of SU-8 2025 (MicroChem, Newton, MA) on a 4-inch silicon wafer (University Wafer, Boston, MA). The wafer was spun at 1800 rpm for 60 s with a ramp of 100 rpm/s in a spin coater (WS-650-23NPP, Laurell Technologies, North Wales, PA) to obtain features of 16 μm height. The wafer was then pre-baked on a hot plate at 65°C for 1 min and at 95°C for 6 min and placed in a table-top laser direct imaging (LDI) system (LPKF ProtoLaser LDI, Garbsen, Germany). The LDI system exposes the structures specified in the software directly to UV light, causing the photoresist to crosslink and solidify. Following the exposure, the wafer was post-baked for 1 min at 65°C and for 6 min at 95°C. By rinsing the wafer with propylene glycol monomethyl ether acetate (PGMEA), the unexposed photoresist was flushed away leaving the desired structures imprinted on the substrate. Finally, the wafer was hard baked for 15 min at 120°C.

The silicon wafer was then used as a mold to fabricate the microfluidic devices. A 9:1 ratio mixture of liquid elastomer (Sylgard 184, Dow Corning, Midland, MI) and curing agent was desiccated to remove air bubbles and cast into the mold. After curing for 60 min at 60°C, the PDMS was removed from the mold. Biopsy punches (0.7 mm diameter, World Precision Instruments, Hitchin, UK) were used to cut fluid access ports into the chip at the position of the inlets. Larger biopsy punches (4 mm diameter, World Precision Instruments, Hitchin, UK) were used to cut the outlet reservoir. The PDMS chip was then plasma-bonded to PDMS-coated coverslips using a standard plasma bonding protocol (100 W, 10 s exposure, 25 sccm, plasma oven from Diener Electric, Ebhausen, Germany).

A crucial step of the OLA protocol is to render the surface of the outlet channel hydrophilic while keeping the LO and IA channel unaltered (14, 15). This was achieved by flushing the outlet channel with a polyvinyl alcohol (PVA) solution for 15 min (50 mg/mL, 87-90% hydrolyzed molecular weight 30,000-70,000 Da, Sigma-Aldrich, St. Louis, MO) via the outer aqueous inlet while applying air pressure from the other inlets. The PVA was removed from the chip by applying suction with a vacuum pump (Gardner Denver Thomas GmbH, Memmingen, Germany). Finally, the microfluidic device was baked in an oven at 120°C for 15 min.

### 2.2 Solution for the Lipid Composition Experiments

A mixture of 200 mM sucrose and 15% v/v glycerol in PBS buffer was used as the standard solution for the inner aqueous (IA) phase of the lipid mixture experiments. The base solution of the outer aqueous (OA) phase was identical to the IA but contained an additional 50 mg/mL poloxamer Kolliphor P-188. For all experiments containing DOPE lipid, P-188 was also added to the IA phase, as we found this increased liposome stability. Furthermore, a solution with the same composition as the IA, but containing 200 mM glucose instead of sucrose was prepared.

Three different binary lipid mixtures were used in the LO phase to test for the membrane composition. The tested lipids were combinations of 1,2-dioleoyl-sn-glycero-3-phospho-rac-(1-glycerol)sodium salt (DOPG), 1,2-dioleoyl-sn-glycero-3-phosphocholine (DOPC) and 1,2-dioleoylsn-glycero-3-phosphoethanolamine (DOPE). The lipids were sourced from Sigma-Aldrich. Aliquots of the individual lipids were combined to form binary lipid systems in three different ratios. The lipids were dissolved in 1-octanol to a final concentration of 3.6 mg/mL to form the LO phase. For each lipid system, one of the lipids of the binary mixture contained a small proportion of a fluorescently labelled lipid (18:1-12:0 NBD PC or 16:1 Liss Rhod PE) which was used to quantify the proportion of the corresponding lipid in the mixture. The exact lipid mixing protocols can be found in the Supplementary Information. The investigated binary lipid systems were as shown in Table 1.

**Table 1:**
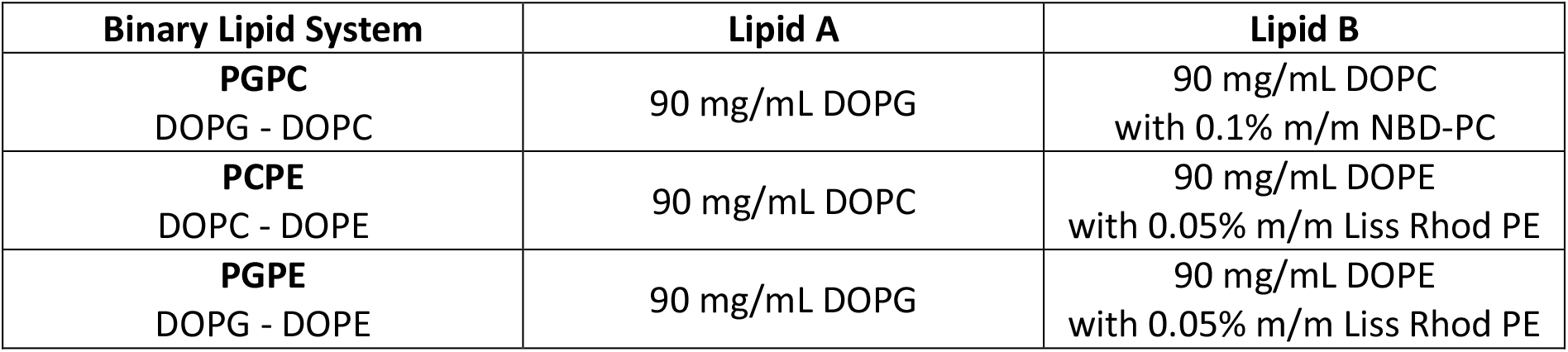
Lipid stocks forming the binary lipid systems used to create GUVs with the OLA technique. Lipid A and Lipid B were combined in 1:3, 2:2, and 3:1 volume ratios each. Lipid B contains a small fraction of fluorescently labelled lipids. The membrane composition was evaluated by observing if the fluorescence intensity of the liposomes scales as expected from the lipid mixture.

### 2.3 Solutions for FRAP Experiments

We formed GUVs of two different PC lipid types by both electroformation and OLA to obtain and compare their lateral diffusion coefficients. The tested lipids were 1,2-dioleoyl-sn-glycero-3-phosphocholine (DOPC) and 1-palmitoyl-2-oleoyl-glycero-3-phosphocholine (POPC). The LO phase consisted of 4 mg/mL PC lipid with 0.5% m/m NBD-PC in octanol. The aqueous solutions (200 mM sucrose, 15% glycerol) were prepared in milli-Q water and not PBS, as the formation of GUVs using electroformation fails at high salt concentrations (13). Two sets of vesicles for each technique were investigated. The lateral diffusion coefficient of one set was measured in a high P-188 environment, the other set was measured in a low P-188 environment. In the high P-188 environment, 50 mg/mL P-188 was encapsulated in the interior of the vesicles, and 14 mg/mL P-188 was present in the surrounding medium for both OLA and electroformed vesicles. The low P-188 environment for electroformation was completely devoid of P-188 (inside and outside the vesicle), while the OLA low P-188 environment had no P-188 encapsulated, but 14 mg/mL P-188 in the surrounding medium. Since OLA requires the addition of P-188 at least in the OA phase to form vesicles, we were not able to create an environment for OLA vesicles that was completely devoid of the poloxamer. The exact solution compositions used for liposome formation and FRAP measurements are listed in Supplementary Tables S1 and S2. We furthermore performed FRAP measurements on electroformed DOPC vesicles at varying levels of glycerol (0% vs 15% glycerol) and different temperatures (approx. 20°C vs. 37°C). For these experiments, we used similar sucrose solutions, devoid of ions and P-188 as in Supplementary Table S1, but with varying glycerol content.

To facilitate imaging, we mixed the vesicles of all experiments with a low-density dilution stock. The dilution stock was similar to the IA solution (no P-188) of the respective experiments but contained 200 mM glucose instead of 200 mM sucrose. The higher molar weight of the sucrose encapsulated within the vesicles leads to a higher density than the surrounding fluid. This causes the vesicles to sink to the bottom of the chip, where they can be imaged more easily.

All chemicals were obtained from Sigma-Aldrich, unless stated otherwise.

### 2.4 Electroformation Protocol

The GUVs were formed using the Vesicle Prep Pro (Nanion Technologies GmbH, Munich, Germany) using an established electroformation protocol (12). 80 μL of a 5 mg/mL lipid suspension (containing 0.5% NBD-PC) in chloroform was spin coated (660 rpm for 2 min) on an Indium Tin Oxide (ITO) coated glass slide (Visiontek) and desiccated for 60 min to evaporate the solvent. 600 μL of the IA solution (Supplementary Table S1) was added and held in place by a rubber O-ring, and sandwiched by another ITO slide. An A/C voltage was applied via the conducting surfaces of the ITO slides inducing swelling of the lipid film and the formation of vesicles (12). The electroformation process was performed at 37°C and ran through the following protocol: the A/C voltage linearly increased from 0 V to 3.2 V peak-to-peak (p-p) at 10 Hz over a time period of 1 hour. Then the voltage stayed at 3.2 V p-p and 10 Hz for 50 minutes. Finally, the frequency decreased linearly to 4 Hz over a time window of 10 minutes and was held at 4 Hz for another 20 minutes. The vesicle suspension was then removed and stored in an Eppendorf tube.

### 2.5 Vesicle Formation and Extraction

The liquid flows were controlled with a pressure-driven microfluidic pump (MFCS-EZ, Fluigent, Le Kremlin-Bicêtre, France) equipped with a Fluiwell-4C reservoir kit. Polymer tubes (Micrewtube 0.5 mL, Simport, Saint-Mathieu-de-Beloeil, Canada) containing the OLA solutions were screwed into the Fluiwell-4C. The solutions entered the microfluidic chip via tygon tubing (microbore tubing, 0.020’’ x 0.060’’ OD, Cole Parmer, St Neots, UK). Cut dispensing tips (Gauge 23 blunt end, Intertronics, Kidlington, UK) were used as metal connectors between the tubing and the chip. Liposome formation was performed by adjusting the respective fluid pressures as previously described (15).

In the lipid composition experiments, the outlet reservoir was used to both collect and directly image the created liposomes. After a stable liposome formation was established, 15 μL of the low-density dilution stock was pipetted into the outlet. The higher density of the liquid inside the liposomes compared to the outside solution caused the liposomes to sink to the bottom of the chip which facilitated imaging. Furthermore, this enabled the separation of the vesicles from the octanol droplets, as octanol has a lower density than water. After 2-3 hours of GUV formation, the microfluidic chip was disconnected from the microfluidic pump and imaged on a confocal microscope.

The vesicles for the FRAP experiments were not imaged directly on the microfluidic chip. Instead, the GUVs were extracted from the chip and transferred to a PDMS coated coverslip with an incubation chamber (Grace Bio-Labs FlexWell, Sigma-Aldrich). After GUV formation was established, 15 μL of the OA solution was added to the outlet reservoir of the OLA chip. After 2-3 hours of liposome formation, 20 μL of the GUV suspension was extracted from the microfluidic chip using a wide bore pipette. The vesicle solution was added to the incubation chamber containing 50 μL of the low-density dilution stock. Similarly, 20 μL of the electroformed vesicle solution was added to 50 μL of the low-density dilution stock in a different visualization chamber. The vesicles were left to settle at the bottom of the visualization chamber for 1 hour before imaging.

### 2.6 Microscopy Parameters and Image Analysis

Standard epifluorescence microscopes (Nikon TE 2000U or Olympus IX 73) were used for imaging the microfluidic devices during vesicle production and PVA treatment of the microfluidic chips (19). The recording of the fluorescence data of the lipid mixtures was performed on commercial inverted confocal microscopes. Images were obtained with the focal plane of the microscope set to the center of the vesicles in order to capture the fluorescence at the equator of the vesicles. A Leica TCS SP5 Confocal was used to image PGPC liposomes fluorescently labelled with nitrobenzoxadiazole (NBD), excited by a 488 nm laser. An Olympus FluoView FV1000 Confocal Laser Scanning Microscope was used to image PGPE and PCPE liposomes fluorescently labelled with Liss Rhod PE, which were excited by a 559 nm laser. Importantly, all optical parameters were kept the same for the measurement of each lipid system. The detailed imaging parameters can be found in the Supplementary Information. The mean intensity values of the fluorescent ring were extracted using the open source software ImageJ, as depicted in Supplementary Figure S1.

We performed a linear regression for each lipid system (PGPC, PCPE and PGPE) with the fluorescence intensities of the liposomes on the y-axis and relative concentrations of the fluorescently tagged lipid in the LO phase on the x-axis. The y-intercept for the regression was fixed at zero. We then normalized the fluorescence intensities of each lipid system with the slope of the linear function we obtained from the regression. By normalizing to the slope of the regression, the new values scale directly with the relative concentrations of the fluorescently doped lipid in the mixture. This results in expected values of 1, 2 and 3 for the 3:1, 2:2 and 1:3 (non-fluorescent: fluorescent lipid ratio) systems, which facilitates comparison of the fluorescence intensity ratios.

The FRAP measurements were performed on an Olympus FluoView FV1000 Confocal Laser Scanning Microscope equipped with a cellVivo Incubation System (PeCon GmbH, Erbach, Germany). The field of view was focused on the bottom of a GUV. By adjusting the pinhole diameter, the slice thickness was increased such that the lower part of a GUV was observed as a fluorescent disc. Using the FRAP function of the microscope’s software, a spot of Ø 4 μm was bleached and the fluorescence recovery observed. 8 images were collected pre-bleaching. Bleaching was performed over 0.1 s with 98% laser power and the fluorescence recovery was recorded for 100 frames (2 μs/pixel exposure).

We calculated the fractional fluorescence recovery trace *f_K_*(*t*) for each vesicle, according to the formula below (30, 31):

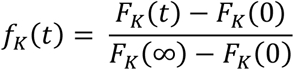

where *F_K_*(*t*) is the measured fluorescence intensity, *F_K_*(0) is the intensity just after bleaching and *F_K_*(∞) is the recovered intensity. The recovered intensity was defined as the average of the last 8 frames of the fluorescence trace. Furthermore, the mobile fraction of each vesicle was calculated:

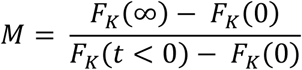

The fluorescence intensity before bleaching *F_K_*(*t* < 0) is defined as the average of the 8 frames recorded pre-bleaching.

An exponential function of the form y = y_0_ (1 − exp(-at)) was fit to the fractional recovery curve of each vesicle and the half-life recovery time *t*_1/2_ was extracted, as shown in Figure 4. We calculated the lipid lateral diffusion coefficient of each vesicle, following the approach of Axelrod *et al*. (30) and Soumpasis (31):

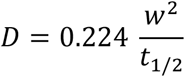

where *w* is the radius of the bleaching spot.

**Figure 2:**
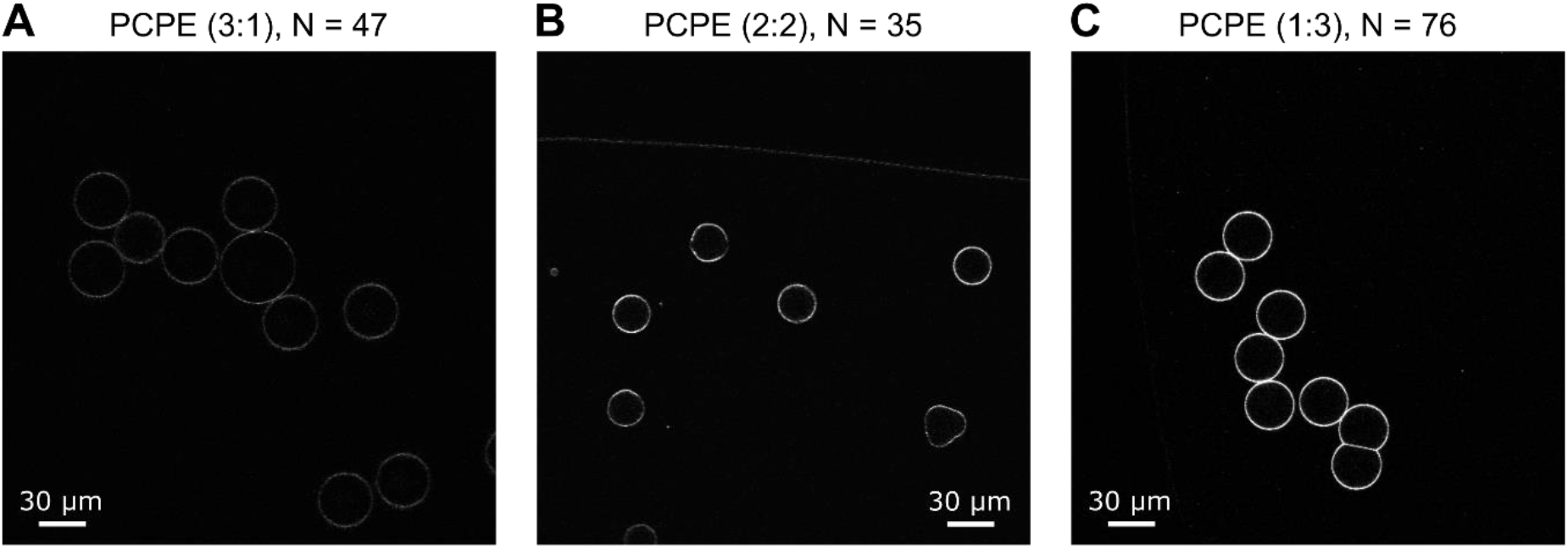
Confocal images of the PCPE (DOPC-DOPE) lipid system in different volume ratios. The fluorescence intensity of the liposomes scales according to the content of fluorescently labelled DOPE in the lipid-octanol phase. PCPE 3:1 (A) vesicles with the least amount of DOPE show the lowest fluorescence intensities, whereas PCPE 1:3 (C) vesicles with the highest content of DOPE in the octanol expresses the strongest fluorescence. PCPE 2:2 (B) with equal amounts of DOPC and DOPE lies in between the two.

**Figure 3:**
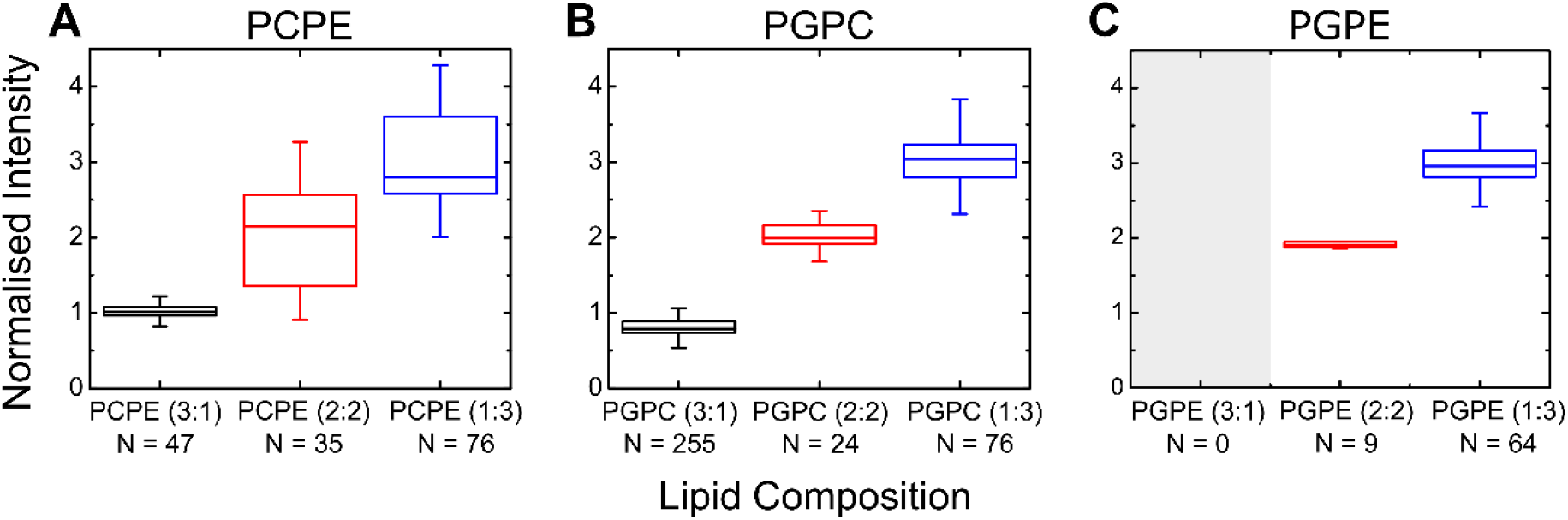
Boxplots of the mean fluorescence intensity analysis for the binary lipid mixtures studied. We normalized the fluorescence intensities to the slope of a linear regression with the fluorescence intensities of the liposomes on the y-axis and relative concentrations of the fluorescently tagged lipid in the LO phase on the x-axis. The normalized intensities of the lipid systems increase in accordance with their larger fraction of the fluorescently doped lipid. The increase in fluorescence for PCPE (A), PGPC (B) and PGPE (C) scales in a linear manner, as expected from the relative concentration of the fluorescently doped lipid in the LO phase. It was not possible to form PGPE lipid vesicles in a 3:1 mixing ratio. The upper and bottom ends of the box indicate the top and bottom quartile, whereas the upper and lower whiskers indicate the smallest and largest value of the set. Outliers ± 3/2 of the upper and lower quartiles are not shown in the plot but are included in the analysis. The line in the middle of the box indicates the median value.

**Figure 4:**
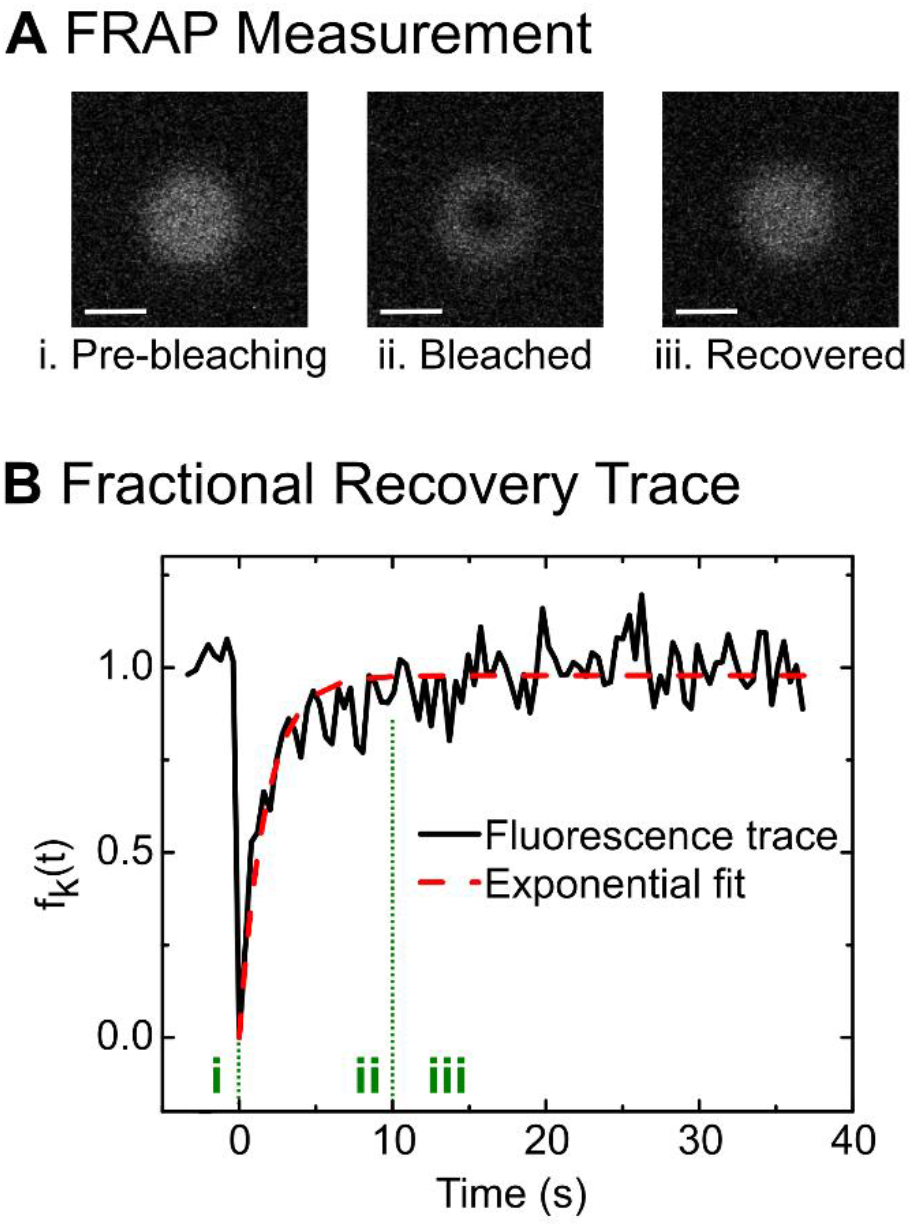
*(A) Example of a vesicle in the different stages of a FRAP measurement. The fluorescence intensity of a circular disk is recorded pre-bleaching (i), bleached (ii) and recovered (iii). The bleached region manifests itself as a dark circle on the vesicle membrane. Scale bar 10 μm. (B) Fractional recovery trace of a vesicle. An exponential curve is fit to the trace from which the halflife recovery time t*_1/2_ *is extracted. The lateral diffusion coefficient of the lipids is calculated using the extracted half-life time and the area of the bleaching spot*.

## 3. Results

### 3.1 Lipid Composition Experiments

We investigated the lipid composition of GUVs formed using the OLA system by performing a mean fluorescence intensity analysis on PGPE, PGPC and PCPE binary lipid mixtures. One of the lipids used contained a small (0.05% or 0.1% m/m) fraction of fluorescently labelled lipids. When forming liposomes in different volume ratios of the two major lipids, the mean fluorescence of the liposomes is expected to scale according to the amount of lipid with the fluorescent label.

Importantly, the images were acquired with identical optical parameters for all three volume ratios of the binary lipid systems. The difference in fluorescence is therefore not the result of a difference in excitation power, but of a higher number of the fluorescently labelled lipids in the GUV membrane. When choosing the microscope parameters, we were careful to eliminate the possibility of PMT saturation, which would have skewed our measurement. We always performed the measurement with the liposomes containing the largest amount of fluorescently labelled lipids first, which was expected to have the highest fluorescence intensity. After calibrating the microscope properties with this set and making sure no PMT saturation occurred, the GUVs with lower expected intensities were imaged.

Figure 2 shows representative images obtained for the lipid mixtures PCPE (DOPC – DOPE) in three different volume ratios. In this case, the DOPE stock contained 0.05% Liss Rhod PE lipids. As can be seen in the images, the fluorescence intensity of the liposomes increases with larger DOPE content in the LO phase, as expected. We also observed this behavior for the other two lipid systems PGPE (DOPG – DOPE) and PGPC (DOPG – DOPC), where the DOPC phase was doped with 0.1% of the fluorescent NBD-PC. Representative images of all three binary lipid systems are shown in the panels of Supplementary Figures S2a, S2b and 2c.

We performed a mean fluorescence intensity analysis on each of the binary lipid systems under investigation in order to quantify the shift in fluorescence between the different lipid mixing ratios. The results are depicted in Figure 3. In the analysis, we performed a linear regression on the fluorescence intensities of each lipid system and then normalized the fluorescence values to the slope of the linear function we obtained. This results in a gradient of +1 for the normalized intensity values with increasing relative concentrations of fluorescently doped lipid. For the 3:1, 2:2 and 1:3 lipid mixtures this translates into values of 1, 2 and 3, respectively, if the lipid composition of the LO phase is maintained in the vesicles produced. We observe the expected 1-2-3 scaling in our experiments. The PCPE vesicles showed values (mean ± std. dev.) of 1.01 ± 0.1, 2.01 ± 0.68 and 2.98 ± 0.62 for the mixing ratios 3:1, 2:2 and 1:3, respectively. The PGPC vesicles yielded mean normalized intensities of 0.81 ± 0.13, 2.06 ± 0.25 and 3.02 ± 0.32 for the mixing ratios 3:1, 2:2 and 1:3, respectively. Note that we were not able to form stable PGPE liposomes in the 3:1 lipid ratio. However, we were able to form PGPE vesicles in the ratios 2:2 and 1:3, which followed the expected scaling with values of 1.99 ± 0.29 and 3.00 ± 0.33.

We attribute the small deviations we observed from a linear increase in intensity to pipetting error, photo bleaching as well as low signal-to-noise ratio. The latter affects primarily the vesicles with low amounts of fluorescent lipid, as we imaged all lipid systems with constant optical parameters and did not change the signal-to-noise ratio by adjusting the gain setting of the microscope.

### 3.2 Lateral Diffusion Measurements

Using the FRAP technique, we measured the lateral diffusion coefficients of lipids in vesicles generated with OLA and compared the results to those obtained in vesicles produced with electroformation. We investigated vesicles of the PC lipid types DOPC and POPC. As stated in section 2.3, we performed the FRAP experiments in two different chemical environments for each technique. OLA requires the use of the Poloxamer P-188 in the OA phase, and we therefore investigated whether or not this had any effect on lipid diffusion in GUV membranes. We performed 4 sets of experiments, 2 each for electroformation and OLA produced GUVs, in varying chemical environments to explore the phase space of possible P-188 combinations. The different chemical compositions of the interior and exterior of the vesicles, along with the production method are reported in Supplementary Table S1 and S2. Furthermore, we investigated the effect of glycerol and temperature on electroformed DOPC vesicles, by performing FRAP on GUVs both in a solution with 0% and 15% glycerol, as well as at room temperature (approx. 20°C) and 37°C, respectively.

We followed the guidelines for FRAP analysis recommended by Chen *et al*. (32) and Tocanne *et al*. (33), only including diffusion measurements performed on vesicles where the radius of the bleached spot *w* was small compared to the diffusion area 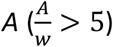. Furthermore, we kept the bleaching pulse *t_B_* short compared to half-life recovery time 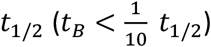 and used *t_B_* = 0.1 *s*, as recommended by Guo *et al*. (34). Additionally, we excluded vesicles that moved during the FRAP measurement, as well as vesicles whose fluorescence did not recover to at least 75% of the pre-bleaching intensity (exclude mobile fraction of M < 0.75%). For the latter, the assumption of an infinite lipid reservoir is not met, and the diffusion coefficient can be underestimated due to the bleaching of substantial parts of the membrane. Typically, these vesicles coincided with the vesicles excluded for one of the other requirements as well.

The lateral lipid diffusion coefficients of DOPC vesicles obtained with the different formation techniques are compared in Figure 5A. Without the presence of P-188 in the IA, the FRAP experiments revealed values (mean ± std. dev.) of 1.0 ± 0.2 μm^2^/s (N = 17) and 1.1 ± 0.2 μm^2^/s (N = 34) for electroformed and OLA vesicles, respectively. Note that in the above case, the outside solution of the OLA vesicles contained 14 mg/mL P-188, whereas the outside solution of the electroformed vesicles was devoid of P-188. As pointed out in section 2.3, the reason for this lies in the fact that GUV formation with OLA is not possible without the presence of P-188 in the OA phase. GUVs formed with 50 mg/mL P-188 encapsulated within the vesicle showed values of 1.2 ± 0.4 μm^2^/s (N = 14) for electroformation and 1.0 ± 0.3 μm^2^/s (N = 30) for OLA. The measurements on POPC vesicles yielded similar results as the DOPC measurements in the range of 1 μm^2^/s. Electroformed and OLA vesicles without the presence of P-188 had lateral diffusion coefficients of 0.8 ± 0.2 μm^2^/s (N = 28) and 1.0 ± 0.3 μm^2^/s (N = 49), respectively. With 50 mg/mL P-188 encapsulated in them, the GUVs yielded diffusion values of 1.3 ± 0.4 μm^2^/s (N = 20) and 0.9 ± 0.3 μm^2^/s (N = 27) for electroformation and OLA, respectively.

**Figure 5:**
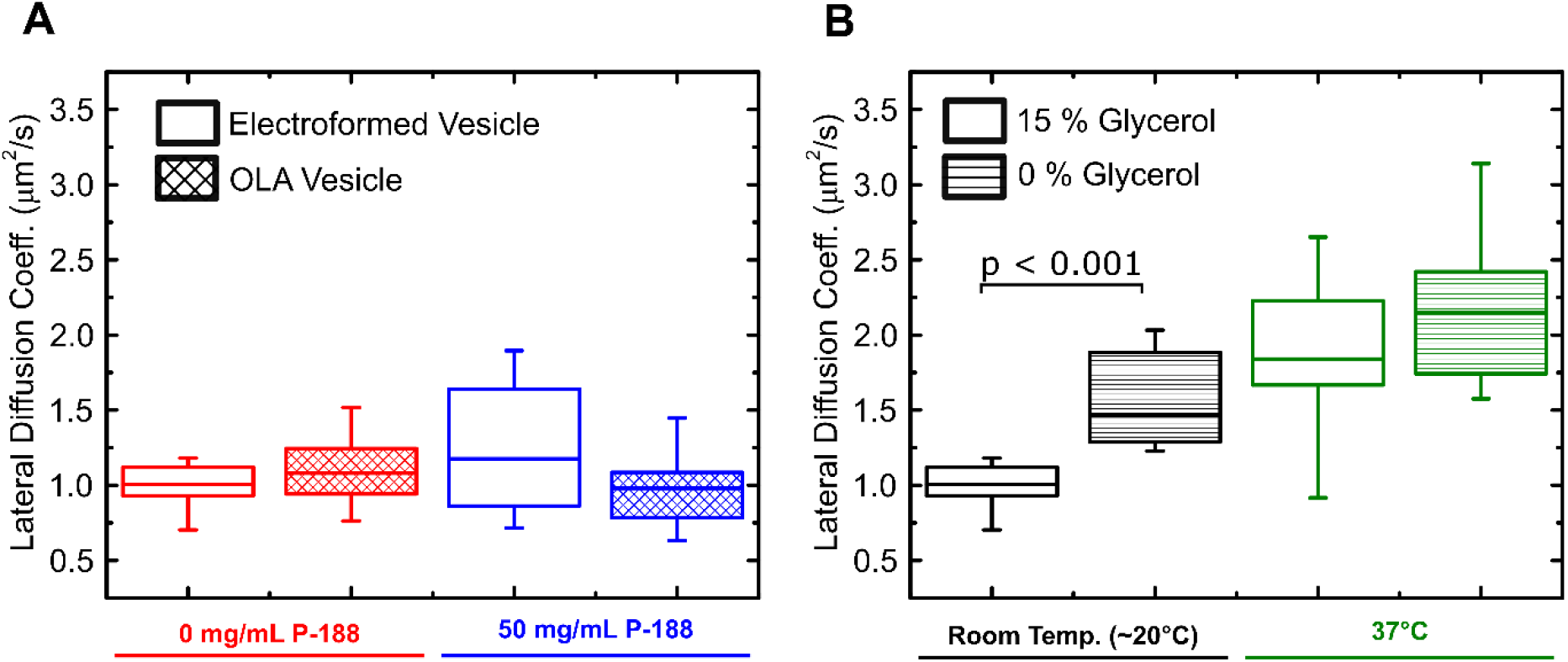
Boxplots of the lipid lateral diffusion coefficients obtained via FRAP. (A) Comparison of DOPC vesicles produced by OLA and electroformation with varying concentrations of encapsulated P-188. The lateral diffusion coefficients are on the order of 1 μm^2^/s for all investigated systems, irrespective of the production method or the presence of P-188. (B) Lipid lateral diffusion coefficients of electroformed DOPC vesicles at varying temperatures and glycerol concentrations. We found a significant (p < 0.001) increase in lateral diffusion with rising temperature and decreasing glycerol concentration, compared to the base line at room temperature (approx. 20°C) and 15% glycerol.

We furthermore conducted FRAP measurements on electroformed DOPC vesicles with varying glycerol content (0% vs. 15%) and temperatures (20°C vs. 37°C), shown in Figure 5B. We found a stronger difference between the lateral diffusion coefficients with varying glycerol and temperatures than between the different formation techniques or varying P-188 concentrations. The diffusion coefficient (mean ± std. dev) increases from 1.0 ± 0.2 μm^2^/s (N = 17) at 20°C and 15% glycerol to 1.6 ± 0.2 μm^2^/s (N = 12) without the presence of glycerol. At 37°C, the coefficients rise to 1.9 ± 0.6 μm^2^/s (N = 19) with 15% glycerol and 2.2 ± 0.5 μm^2^/s (N = 7) without glycerol.

Interestingly, we were able to find vesicles with and without an attached octanol pocket amongst the population of extracted OLA vesicles for both DOPC and POPC. Supplementary Figure S3 shows isometric and confocal sliced views of GUVs with and without the octanol attached. We did not observe a significant (p < 0.01) difference between the lateral diffusion coefficients of vesicles with and without octanol pockets attached. The FRAP measurements of both DOPC and POPC, as well as the measurements with varying temperature and glycerol content are summarized again in Supplementary Tables S3 and S4. Statistical analyses for the FRAP measurements are reported in Supplementary Tables S5-S12.

## 4. Discussion

### 4.1 Lipid Mixture Experiments

Since the introduction of electroformation by Angelova and Dimitrov in 1986 (12), this technique has been widely adopted in the biophysics community for the creation and study of model membranes (35). For instance, the method has been used to investigate the membrane phase behavior (36) and mechanical properties (37) of lipid bilayers. In addition, electroformation has also been used for the creation of liposomes with complex binary and ternary lipid mixtures (36, 38). Our experiments show that OLA is likewise able to form GUVs of different binary lipid mixtures. Furthermore, OLA provides the advantages that are typically associated with microfluidic techniques to obtain GUVs (13). For instance, the GUV formation is not affected by presence of ions or buffers (14), thereby allowing measurements of membrane properties in an environment more closely mimicking physiological conditions. Furthermore, the inside and outside solutions in OLA are separated as of formation, allowing for selective encapsulation of substances inside the GUVs (18).

The issue impeding our measurements with vesicles of PGPE 3:1 lipid ratio lies in the low stability of these GUVs. Although it was initially possible for us to form these vesicles at the OLA junction, they appeared to be less resistant to mechanical stress compared to the PCPE and PGPC vesicles. The vast majority of PGPE (3:1) vesicles that were created at the OLA junction burst as they flowed through the microfluidic chip towards the outlet reservoir. The likely reason is that these vesicles burst when subjected to shear stress from the PDMS channel walls (19). Although occasionally individual vesicles survived to the end of the outlet channel in the reservoir, we noticed bursting events for these vesicles after several minutes as well.

We partially attribute this behavior to the lipid polymorphism of PGPE (3:1) vesicles. PE lipids are known to have a cone like shape which makes it energetically unfavorable for them to form lamellar structures (39). If forced into a GUV forming bilayer, the acyl chains are pressed together, increasing the lateral pressure at the center of the membrane, a state coined ‘frustrated bilayer’ (40, 41). In nature, this pressure can be balanced by enrichment of the non-bilayer lipid in the inner leaflet of the membranes of cells (40, 42). The fact that we could produce PCPE liposomes in all three lipid ratios suggests that other effects in addition to the lipid shape are responsible for the low stability of PGPE (3:1) vesicles. Additionally, there are reports that PG stabilizes PE membranes, which seemingly contradicts our findings (43). However, these studies only looked at PG fractions of up to 30 mol%, whereas the PGPE (3:1) GUVs in our experiments predominantly consist of PG. MD simulations by Murzyn *et al*. on POPG-POPE (1:3) bilayers revealed that the prevailing interactions between lipid molecules are water bridges and H-bonds (44). While PE predominantly forms all these bonds with PG lipids, PE also bonds to other PE molecules. PG on the other hand barely bonds with other PG molecules (44). The low H-bonding capacity of PG lipids has also been observed in MD simulations on pure PG bilayers by Zhao *et al*., who attribute this to the net negative charge and electrostatic repulsion of the individual molecules (45). However, the two simulations diverge in the role of ion bridges between lipids. Whereas Murzyn *et al*. found that Na^+^ ion bridges are only a minor contributor to membrane stability, Zhao *et al*. found strong ion-mediated interactions between the lipid molecules causing attractive forces that overcome the electrostatic repulsion between the negatively charged PG headgroups (45). Our findings suggest that in addition to the effect of PE, the high content of charged PG in the PGPE (3:1) GUVs further destabilizes the membrane. However, more research is needed to explain the low stability behavior of the PGPE (3:1) vesicles that we observed.

### 4.2 Lateral Diffusion Experiments

Lateral lipid diffusion values reported in the literature vary greatly, as these are strongly affected not only by the chemical and physical environment (46), but also by the choice of measurement technique. Different techniques like Fluorescence Correlation Spectroscopy (FCS), FRAP and NMR have yielded different lateral diffusion coefficients (47). For instance, Filippov *et al*. obtained values of 9.32 μm^2^/s for DOPC and 8.87 μm^2^/s for POPC with NMR (48), both higher than the values obtained by us. Furthermore, the choice of membrane platform, e.g. supported lipid bilayer (SLB) vs. GUV (49) can influence the measurement. Guo *et al*. report POPC lateral diffusion coefficients of 1.8 ± 0.2 μm^2^/s on supported lipid bilayers and 3.3 ± 0.2 μm^2^/s on GUVs (34) while Pincet *et al*. measured DOPC lateral diffusion coefficients of 1.9 ± 0.4 μm^2^/s on supported lipid bilayers and 3.4 ± 0.7 μm^2^/s on GUVs (49). These values are in good agreement with the values we obtained for OLA vesicles, albeit still significantly elevated. We explain this by the presence of 15% v/v glycerol in the OLA solution, as both previous research and our control experiments reveal that glycerol lowers the lateral diffusion coefficient (50). Our FRAP measurements at elevated temperatures and without the presence of glycerol match the above mentioned values more closely. Furthermore, differences in the exact shape of the bleaching profiles, the imaging parameters and data analysis can skew the obtained diffusion values (51). Interestingly, our control experiments also reveal that varying glycerol concentration and temperature both have a stronger effect on the lateral diffusion coefficient of electroformed DOPC vesicles than variations in the P-188 concentration or the formation technique.

Within the investigated population of extracted DOPC and POPC OLA vesicles, we found vesicles with a visible octanol pocket attached, as well as those without visible octanol pockets attached. In the confocal scan of a vesicle, the octanol pocket manifests itself as a bright spot at the top side of a GUV. This location is plausible, due to the lower density of octanol compared to the surrounding aqueous solution. The brightness of the pocket compared to the rest of the vesicle suggests a high amount of lipid dissolved in the octanol. 3D reconstructions of confocal scans of both types of vesicles are depicted in Supplementary Figure S3. Remarkably, we found no significant differences (p < 0.01) in lateral lipid diffusion coefficients between vesicles with or without an octanol pocket. This is an interesting parallel to research by Karamdad *et al*. who compared the bending rigidity of vesicles obtained with a microfluidic technique (using squalene oil) to electroformed vesicles. They also found that the presence of residual oil in the membrane did not alter the biophysical properties of the membrane (52).

While most studies report octanol having a lowering effect on the phase transition temperature of lipids (53–55), conflicting reports exist on its effect on lateral lipid diffusion coefficients. Molecular Dynamics simulations by Griepernau *et al*. on DMPC membranes showed a decrease in lateral diffusion of the lipids in presence of 1-octanol (54), whereas NMR experiments by Rifici *et al*. on the same lipid revealed an increase in lateral diffusion (55). They furthermore reported sudden changes in lateral diffusion near the phase transition temperature T_m_ (55). A possible explanation for why we did not observe a strong shift in lateral diffusion coefficients between OLA vesicles containing octanol vs. electroformed vesicles without octanol might lie in the phase transition temperature of the lipids used in our experiment. According to the manufacturer, DMPC lipids investigated by the previously mentioned groups have a melting temperature of 24°C, whereas the DOPC and POPC lipids we investigated have melting temperatures of −17°C and −2°C, respectively. Since we performed our experiments at room temperature (approximately 20°C), the lipids are well in the fluid phase and the effect of altered phase transition temperature due to the octanol might have diminished. Furthermore, it is possible that the diffusion lowering effect of the glycerol dominates our system. In addition, the so-called cutoff effect of anesthetics could play a role in modulating membrane properties. The cutoff effect describes the phenomenon of the increasing anesthetic potency of alcohols with increasing chain length, which suddenly levels off and even reverses for much longer chain lengths (54–56). Since the investigated DOPC and POPC lipids have 18:1c9 and 16:0-18:1 acyl chains, respectively, octanol with a chain length of 8 might reach into the domain where membrane modulation changes from a destabilizing to a stabilizing effect. However, in this case, the effect should have also been observed in the 14:0 DMPC system.

Overall, our experiments suggest that neither the OLA formation technique, nor the presence of P-188 changes the lateral diffusion coefficients in a substantial manner (see Supplementary Table S3) excluding glycerol or temperature effects. Future experiments to characterize OLA produced vesicles could involve the quantification of the actual content of octanol left in the membrane (if any) after the budding off process, for instance by mass or Raman spectroscopy. A follow up investigation would involve OLA vesicles of different lipid types with higher chain melting temperatures, as they are likely to show more pronounced changes in membrane properties due to the presence of any residual octanol.

## 5. Conclusion

In this work, we showed that the lipid composition of vesicles formed with the novel Octanol-Assisted Liposome Assembly (OLA) technique matches the composition of the lipid in the LO phase input during the vesicle formation process. We did not observe any demixing effects, or cases with one element preferentially remaining in the octanol phase upon liposome production, showing that the technique reliably produces vesicles of desired lipid compositions. In addition, our lipid composition experiments revealed the stable vesicle production of binary lipid mixtures of DOPG-DOPC as well was DOPC-DOPE in 1:3, 2:2 and 3:1 ratios. However, DOPG-DOPE lipid mixtures could only be formed in 2:2 and 3:1 ratios. We hypothesize that the low stability of PGPE vesicles with high (>50%) PG content is due to the polymorphism and charge density of the PE and PG lipids, respectively.

Future experiments involving the use of lysolipids could provide further evidence to indicate whether the cone shape of PE lipids and the charge density of PG are responsible for the low stability of the PGPE (3:1) vesicles we observed. Lysolipids, such as LPC, only have one acyl chain and add a high positive curvature to the membrane. As such, they can counterbalance the negative curvature induced by the PE lipids (39, 40). A tertiary lipid mixture of LPC, DOPG and DOPE should therefore have a higher stability than binary PGPE mixtures. Other potential methods to yield GUVs of arbitrary compositions involve stabilizing the membrane mechanically using nanostructures. Since OLA allows for the efficient encapsulation of substances in the interior of the vesicles, as well as coating from the exterior in well-defined conditions (18), an artificial cytoskeleton could be applied to the membrane, such as a DNA cytoskeleton (57).

Furthermore, we compared the lateral lipid diffusion coefficients of DOPC and POPC liposomes generated using both OLA and electroformation. We found the lateral diffusion coefficients for DOPC and POPC vesicles to be on the order of 1 μm^2^/s for all the chemical compositions and formation protocols studied. The lateral lipid diffusion coefficients of the vesicles generated by the two techniques, OLA and electroformation, showed relatively minor deviations from one another, and additionally most of these differences were found to be statistically insignificant (SI Tables S5-S12). In contrast, an increase in temperature and the removal of glycerol from the vesicle solution resulted in a more than two fold increase in the lateral lipid diffusion coefficient (p < 0.001). Moreover, we were able to compare the lateral lipid diffusion coefficients of OLA vesicles with and without octanol pockets and found that the octanol pocket does not alter the lateral diffusion properties in a statistically significant manner (at the p < 0.01 significance level). Overall, this set of biophysical characterizations demonstrates the similarities in membrane properties for vesicles produced using OLA and electroformation, suggesting that the added functionality of the OLA platform does not involve any compromise in membrane quality. We envisage OLA as being a game changer for the production and study of biomimetic membranes, with major advantages over traditional vesicle formation techniques, for use in synthetic biology, drug testing and the study of membrane biophysics.

## 6. Acknowledgements

M.S acknowledges funding from the Friedrich-Naumann-Foundation and the Jane Bourque-Driscoll Fund. D.S. acknowledges funding from Winton Programme for the Physics of Sustainability and Engineering and Physical Sciences Research Council (EPSRC). J.C. acknowledges funding from a Wellcome Trust Institutional Strategic Support Award to the University of Exeter (204909/Z/16/Z). UFK was supported by an ERC Consolidator Grant (Designer-Pores 647144). We would furthermore like to acknowledge S. Deshpande and K. Al Nahas for valuable discussions and feedback.

## References

1. Deamer, D.W. 2010. From “Banghasomes” to liposomes: A memoir of Alec Bangham, 1921–2010. FASEB J. 24: 1308–1310.

2. Bangham, A.D., and R.W. Horne. 1964. Negative staining of phospholipids and their structural modification by surface-active agents as observed in the electron microscope. J. Mol. Biol. 8: 660–IN10.

3. Akbarzadeh, A., R. Rezaei-Sadabady, S. Davaran, S.W. Joo, N. Zarghami, Y. Hanifehpour, M. Samiei, M. Kouhi, and K. Nejati-Koshki. 2013. Liposome: classification, preparation, and applications. Nanoscale Res. Lett. 8: 102.

4. Göpfrich, K., I. Platzman, and J.P. Spatz. 2018. Mastering Complexity: Towards Bottom-up Construction of Multifunctional Eukaryotic Synthetic Cells. Trends Biotechnol. 36: 938–951.

5. Xu, C., S. Hu, and X. Chen. 2016. Artificial cells: from basic science to applications. Mater. Today. 19: 516–532.

6. Cama, J., C. Chimerel, S. Pagliara, A. Javer, and U.F. Keyser. 2014. A label-free microfluidic assay to quantitatively study antibiotic diffusion through lipid membranes. Lab Chip. 14: 2303–2308.

7. Cama, J., M. Schaich, K. Al Nahas, S. Hernández-Ainsa, S. Pagliara, and U.F. Keyser. 2016. Direct Optofluidic Measurement of the Lipid Permeability of Fluoroquinolones. Sci. Rep. 6: 32824.

8. Cama, J., H. Bajaj, S. Pagliara, T. Maier, Y. Braun, M. Winterhalter, and U.F. Keyser. 2015. Quantification of Fluoroquinolone Uptake through the Outer Membrane Channel OmpF of Escherichia coli. J. Am. Chem. Soc. 137: 13836–13843.

9. Shiomi, H., S. Tsuda, H. Suzuki, and T. Yomo. 2014. Liposome-Based Liquid Handling Platform Featuring Addition, Mixing, and Aliquoting of Femtoliter Volumes. PLoS One. 9: e101820.

10. Walde, P., K. Cosentino, H. Engel, and P. Stano. 2010. Giant Vesicles: Preparations and Applications. ChemBioChem. 11: 848–865.

11. Dimova, R. 2019. Giant Vesicles and Their Use in Assays for Assessing Membrane Phase State, Curvature, Mechanics, and Electrical Properties. Annu. Rev. Biophys. 48: 93–119.

12. Angelova, M.I., and D.S. Dimitrov. 1986. Liposome electroformation. Faraday Discuss. Chem. Soc. 81: 303.

13. van Swaay, D., and A. DeMello. 2013. Microfluidic methods for forming liposomes. Lab Chip. 13: 752.

14. Deshpande, S., Y. Caspi, A.E.C. Meijering, and C. Dekker. 2016. Octanol-assisted liposome assembly on chip. Nat. Commun. 7: 10447.

15. Deshpande, S., and C. Dekker. 2018. On-chip microfluidic production of cell-sized liposomes. Nat. Protoc. 13: 856–874.

16. Teh, S.-Y., R. Khnouf, H. Fan, and A.P. Lee. 2011. Stable, biocompatible lipid vesicle generation by solvent extraction-based droplet microfluidics. Biomicrofluidics. 5: 044113.

17. Neužil, P., S. Giselbrecht, K. Länge, T.J. Huang, and A. Manz. 2012. Revisiting lab-on-a-chip technology for drug discovery. Nat. Rev. Drug Discov. 11: 620–632.

18. Al Nahas, K., J. Cama, M. Schaich, K. Hammond, S. Deshpande, C. Dekker, M.G. Ryadnov, and U.F. Keyser. 2019. A microfluidic platform for the characterisation of membrane active antimicrobials. Lab Chip. 19: 837–844.

19. Schaich, M., J. Cama, K. Al Nahas, D. Sobota, H. Sleath, K. Jahnke, S. Deshpande, C. Dekker, and U.F. Keyser. 2019. An Integrated Microfluidic Platform for Quantifying Drug Permeation across Biomimetic Vesicle Membranes. Mol. Pharm. 16: 2494–2501.

20. Deshpande, S., W.K. Spoelstra, M. van Doorn, J. Kerssemakers, and C. Dekker. 2018. Mechanical Division of Cell-Sized Liposomes. ACS Nano. 12: 2560–2568.

21. Deshpande, S., S. Wunnava, D. Hueting, and C. Dekker. 2019. Membrane Tension– Mediated Growth of Liposomes. Small. 15: 1902898.

22. Deshpande, S., F. Brandenburg, A. Lau, M.G.F. Last, W.K. Spoelstra, L. Reese, S. Wunnava, M. Dogterom, and C. Dekker. 2019. Spatiotemporal control of coacervate formation within liposomes. Nat. Commun. 10: 1800.

23. Deshpande, S., A. Birnie, and C. Dekker. 2017. On-chip density-based purification of liposomes. Biomicrofluidics. 11: 034106.

24. Rowe, E.S., F. Zhang, T.W. Leung, J.S. Parr, and P.T. Guy. 1998. Thermodynamics of Membrane Partitioning for a Series of n-Alcohols Determined by Titration Calorimetry: Role of Hydrophobic Effects. Biochemistry. 37: 2430–2440.

25. Cantor, R.S. 2001. Bilayer Partition Coefficients of Alkanols: Predicted Effects of Varying Lipid Composition. J. Phys. Chem. B. 105: 7550–7553.

26. Duncan, A.L., T. Reddy, H. Koldsø, J. Hélie, P.W. Fowler, M. Chavent, and M.S.P. Sansom. 2017. Protein crowding and lipid complexity influence the nanoscale dynamic organization of ion channels in cell membranes. Sci. Rep. 7: 16647.

27. Zhang, F., G.M. Lee, and K. Jacobson. 1993. Protein lateral mobility as a reflection of membrane microstructure. BioEssays. 15: 579–588.

28. Jacobson, K., P. Liu, and B.C. Lagerholm. 2019. The Lateral Organization and Mobility of Plasma Membrane Components. Cell. 177: 806–819.

29. Yang, K., and X. Han. 2016. Lipidomics: Techniques, Applications, and Outcomes Related to Biomedical Sciences. Trends Biochem. Sci. 41: 954–969.

30. Axelrod, D., D.E. Koppel, J. Schlessinger, E. Elson, and W.W. Webb. 1976. Mobility measurement by analysis of fluorescence photobleaching recovery kinetics. Biophys. J. 16: 1055–1069.

31. Soumpasis, D.M. 1983. Theoretical analysis of fluorescence photobleaching recovery experiments. Biophys. J. 41: 95–97.

32. Chen, Y., B.C. Lagerholm, B. Yang, and K. Jacobson. 2006. Methods to measure the lateral diffusion of membrane lipids and proteins. Methods. 39: 147–153.

33. Tocanne, J.-F., L. Dupou-Cézanne, and A. Lopez. 1994. Lateral diffusion of lipids in model and natural membranes. Prog. Lipid Res. 33: 203–237.

34. Guo, L., J.Y. Har, J. Sankaran, Y. Hong, B. Kannan, and T. Wohland. 2008. Molecular Diffusion Measurement in Lipid Bilayers over Wide Concentration Ranges: A Comparative Study. ChemPhysChem. 9: 721–728.

35. Beales, P.A., B. Ciani, and A.J. Cleasby. 2015. Nature’s lessons in design: nanomachines to scaffold, remodel and shape membrane compartments. Phys. Chem. Chem. Phys. 17: 15489–15507.

36. Veatch, S.L., and S.L. Keller. 2005. Seeing spots: Complex phase behavior in simple membranes. Biochim. Biophys. Acta - Mol. Cell Res. 1746: 172–185.

37. Faizi, H.A., S.L. Frey, J. Steinkühler, R. Dimova, and P.M. Vlahovska. 2019. Bending rigidity of charged lipid bilayer membranes. Soft Matter. 15: 6006–6013.

38. Korlach, J., P. Schwille, W.W. Webb, and G.W. Feigenson. 1999. Characterization of lipid bilayer phases by confocal microscopy and fluorescence correlation spectroscopy. Proc. Natl. Acad. Sci. 96: 8461–8466.

39. Frolov, V.A., A. V. Shnyrova, and J. Zimmerberg. 2011. Lipid Polymorphisms and Membrane Shape. Cold Spring Harb. Perspect. Biol. 3: a004747–a004747.

40. van den Brink-van der Laan, E., J. Antoinette Killian, and B. de Kruijff. 2004. Nonbilayer lipids affect peripheral and integral membrane proteins via changes in the lateral pressure profile. Biochim. Biophys. Acta - Biomembr. 1666: 275–288.

41. de Kruijff, B. 1997. Lipid polymorphism and biomembrane function. Curr. Opin. Chem. Biol. 1: 564–9.

42. Devaux, P.F. 1991. Static and dynamic lipid asymmetry in cell membranes. Biochemistry. 30: 1163–1173.

43. Tari, A., and L. Huang. 1989. Structure and function relationship of phosphatidylglycerol in the stabilization of phosphatidylethanolamine bilayer. Biochemistry. 28: 7708–7712.

44. Murzyn, K., T. Róg, and M. Pasenkiewicz-Gierula. 2005. Phosphatidylethanolamine-Phosphatidylglycerol Bilayer as a Model of the Inner Bacterial Membrane. Biophys. J. 88: 1091–1103.

45. Zhao, W., T. Róg, A.A. Gurtovenko, I. Vattulainen, and M. Karttunen. 2007. Atomic-Scale Structure and Electrostatics of Anionic Palmitoyloleoylphosphatidylglycerol Lipid Bilayers with Na+ Counterions. Biophys. J. 92: 1114–1124.

46. Almeida, P.F.F., W.L.C. Vaz, and T.E. Thompson. 1992. Lateral diffusion in the liquid phases of dimyristoylphosphatidylcholine/cholesterol lipid bilayers: a free volume analysis. Biochemistry. 31: 6739–6747.

47. Macháň, R., Y.H. Foo, and T. Wohland. 2016. On the Equivalence of FCS and FRAP: Simultaneous Lipid Membrane Measurements. Biophys. J. 111: 152–161.

48. Filippov, A., G. Orädd, and G. Lindblom. 2003. Influence of Cholesterol and Water Content on Phospholipid Lateral Diffusion in Bilayers. Langmuir. 19: 6397–6400.

49. Pincet, F., V. Adrien, R. Yang, J. Delacotte, J.E. Rothman, W. Urbach, and D. Tareste. 2016. FRAP to Characterize Molecular Diffusion and Interaction in Various Membrane Environments. PLoS One. 11: e0158457.

50. Orädd, G., G. Wikander, G. Lindblom, and L.B.-Å. Johansson. 1994. Effect of glycerol on the translational and rotational motions in lipid bilayers studied by pulsed-field gradient 1 H NMR, EPR and time-resolved fluorescence spectroscopy. J. Chem. Soc., Faraday Trans. 90: 305–309.

51. Jönsson, P., M.P. Jonsson, J.O. Tegenfeldt, and F. Höök. 2008. A Method Improving the Accuracy of Fluorescence Recovery after Photobleaching Analysis. Biophys. J. 95: 5334–5348.

52. Karamdad, K., R. V. Law, J.M. Seddon, N.J. Brooks, and O. Ces. 2015. Preparation and mechanical characterisation of giant unilamellar vesicles by a microfluidic method. Lab Chip. 15: 557–562.

53. Blicher, A., K. Wodzinska, M. Fidorra, M. Winterhalter, and T. Heimburg. 2009. The Temperature Dependence of Lipid Membrane Permeability, its Quantized Nature, and the Influence of Anesthetics. Biophys. J. 96: 4581–4591.

54. Griepernau, B., S. Leis, M.F. Schneider, M. Sikor, D. Steppich, and R.A. Böckmann. 2007. 1-Alkanols and membranes: A story of attraction. Biochim. Biophys. Acta - Biomembr. 1768: 2899–2913.

55. Rifici, S., G. D’Angelo, C. Crupi, C. Branca, V. Conti Nibali, C. Corsaro, and U. Wanderlingh. 2016. Influence of Alcohols on the Lateral Diffusion in Phospholipid Membranes. J. Phys. Chem. B. 120: 1285–1290.

56. Ingólfsson, H.I., and O.S. Andersen. 2011. Alcohol’s Effects on Lipid Bilayer Properties. Biophys. J. 101: 847–855.

57. Kurokawa, C., K. Fujiwara, M. Morita, I. Kawamata, Y. Kawagishi, A. Sakai, Y. Murayama, S.M. Nomura, S. Murata, M. Takinoue, and M. Yanagisawa. 2017. DNA cytoskeleton for stabilizing artificial cells. Proc. Natl. Acad. Sci. 114: 7228–7233.

