## Supplementary Information for "Biophysical characterization reveals the similarities of liposomes produced using microfluidics and electroformation"

### 1. Lipid Composition Experiments

#### 1.1. Preparation of Lipid Stocks

The binary lipid mixtures for the liposome composition experiments were obtained as follows:

- **PGPC (DOPG – DOPC):** DOPC and DOPG lipid powder was dissolved in 100% ethanol to a final concentration of 100 mg/mL in separate vials. Furthermore, 18:1-12:0 NBD-PC was dissolved in ethanol to a final concentration of 1 mg/mL. An aliquot of fluorescently doped DOPC lipid was made by mixing the DOPC and NBD-PC in a 9:1 ratio, resulting in a total concentration of 90 mg/mL DOPC lipid. Similarly, a DOPG aliquot was obtained by mixing it in a 9:1 ratio with pure ethanol, resulting in a total concentration of 90 mg/mL DOPG lipid. The 90 mg/mL DOPG and DOPC/NBD-PC aliquots were further mixed in 1:3, 2:2 and 3:1 volume ratios and used as lipid stocks for the OLA experiments. The lipid stocks were stored in the freezer at -20°C. The binary lipid stocks were further diluted to 3.6 mg/mL in 1-octanol before each experiment.

- **PCPE (DOPC - DOPE):** DOPC and DOPE lipid powder was dissolved in 100% ethanol to a final concentration of 100 mg/mL in separate vials. Furthermore, 16:0 Liss Rhod PE was dissolved in ethanol to a final concentration of 0.5 mg/mL. An aliquot of fluorescently doped DOPE lipid was made by mixing the DOPE and Liss Rhod PE in a 9:1 ratio, resulting in a total concentration of 90 mg/mL DOPE lipid. Similarly, a DOPC aliquot was obtained by mixing it in a 9:1 ratio with pure ethanol, resulting in a total concentration of 90 mg/mL DOPC lipid. The 90 mg/mL DOPC and DOPE/Liss Rhod PE aliquots were further mixed in 1:3, 2:2 and 3:1 volume ratios and used as lipid stocks for the OLA experiments. The lipid stocks were stored in the freezer at -20°C. The binary lipid stocks were further diluted to 3.6 mg/mL in 1-octanol before each experiment.

- **PGPE (DOPG - DOPE):** DOPG and DOPE lipid powder was dissolved in 100% ethanol to a final concentration of 100 mg/mL in separate vials. Furthermore, 16:0 Liss Rhod PE was dissolved in ethanol to a final concentration of 0.5 mg/mL. An aliquot of fluorescently doped DOPE lipid was made by mixing the DOPE and Liss Rhod PE in a 9:1 ratio, resulting in a total concentration of 90 mg/mL DOPE lipid. Similarly, a DOPC aliquot was obtained by mixing it in a 9:1 ratio with pure ethanol, resulting in a total concentration of 90 mg/mL DOPG lipid. The 90 mg/mL DOPG and DOPE/Liss Rhod PE aliquots were further mixed in 1:3, 2:2 and 3:1 volume ratios and used as lipid stocks for the OLA experiments. The lipid stocks were stored in the freezer at -20°C. The binary lipid stocks were further diluted to 3.6 mg/mL in 1-octanol before each experiment.

All chemicals were acquired from Sigma-Aldrich, unless stated otherwise.

### 1.2.Optical Parameters

The confocal images were obtained using the following optical parameters:

- **PGPC (DOPG – DOPC):** The images were acquired on a Leica TCS SP5 confocal microscope, equipped with a 488 nm Argon laser. The laser output power was set to 10%. Images were recorded via a 40× oil immersion objective (HCX PL APO CS 40.0, NA 1.25) with a scan speed of 400 Hz. Pinhole Diameter 67.93 µm, Laser Acousto-Optic Tunable Filter (AOTF) 40%, Smart Gain (HyD) 302% and Smart Offset disabled. The microscope was controlled via the Leica Microsystems LAS AF software.

- **PGPE (DOPG – DOPE):** The images were acquired in an Olympus FV 1000 confocal microscope, equipped with a LD559 laser (559 nm, 15 mW). The laser output power was set to 1%. Images were recorded via a 40× oil immersion objective (UPFLN 40×, NA 1.3) with a scan speed of 2 µs/pixel. Pinhole diameter 80 µm, PMT offset voltage 645 V and analog PMT offset 0. The microscope was controlled via the Olympus microscope Fluoview software.

- **PCPE (DOPC – DOPE):** The images were acquired in an Olympus FV 1000 confocal microscope, equipped with a LD559 laser (559 nm, 15 mW). The laser output power was set to 1%. Images were recorded via a 40× oil immersion objective (UPFLN 40×, NA 1.3) with a scan speed of 2 µs/pixel. Pinhole diameter 80 µm, PMT offset voltage 645 V and analog PMT offset 0. The microscope was controlled via the Olympus microscope Fluoview software.

### 1.3.Image Analysis

Image analysis was performed using the open source software ImageJ, as depicted in Figure S1. Using the software's band tool, the intensity of the fluorescent ring was extracted. The band width was 2 µm. Overlaps with other vesicles, or bright fluorescent spots stemming from residue octanol in the membrane were excluded from the analysis using the software's brush tool.

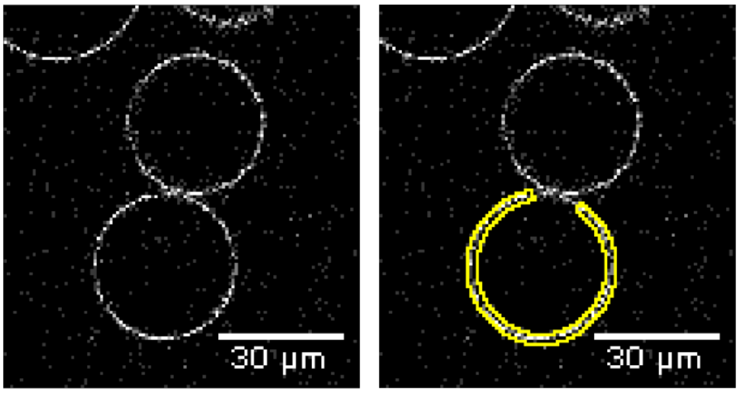


***Figure S1:*** *Extraction of the mean average fluorescence of GUVs. A 2 µm thick band (shown in yellow) engulfing the fluorescent ring of the liposome was used as the region of interest. Areas of overlapping liposomes, or bright fluorescent spots stemming from residual octanol or lipid aggregates in the membrane were excluded.*

#### 1.4. Image Panels of Binary Lipid Systems

Representative images of GUVs obtained for the different binary lipid systems are shown in Figures S2a, S2b and S2c, along with boxplots of their corresponding mean fluorescence intensity analyses. A clear increase in intensity of the vesicles for the different lipid mixtures can be observed. In the analysis, we performed a linear regression on the fluorescence intensities of each lipid system and then normalized the fluorescence values to the slope of the linear function we obtained. This results in a gradient of +1 for the normalized intensity values with increasing relative concentrations of fluorescently doped lipid. For the 3:1, 2:2 and 1:3 lipid mixtures this translates into values of 1, 2 and 3, respectively, if the lipid composition of the LO phase is maintained in the vesicle.

For the PGPC and PCPE systems, the normalized intensity shows a clear 1-2-3 increase, as expected. Stable PGPE vesicles could only be formed in the 2:2 and 1:3 ratio.

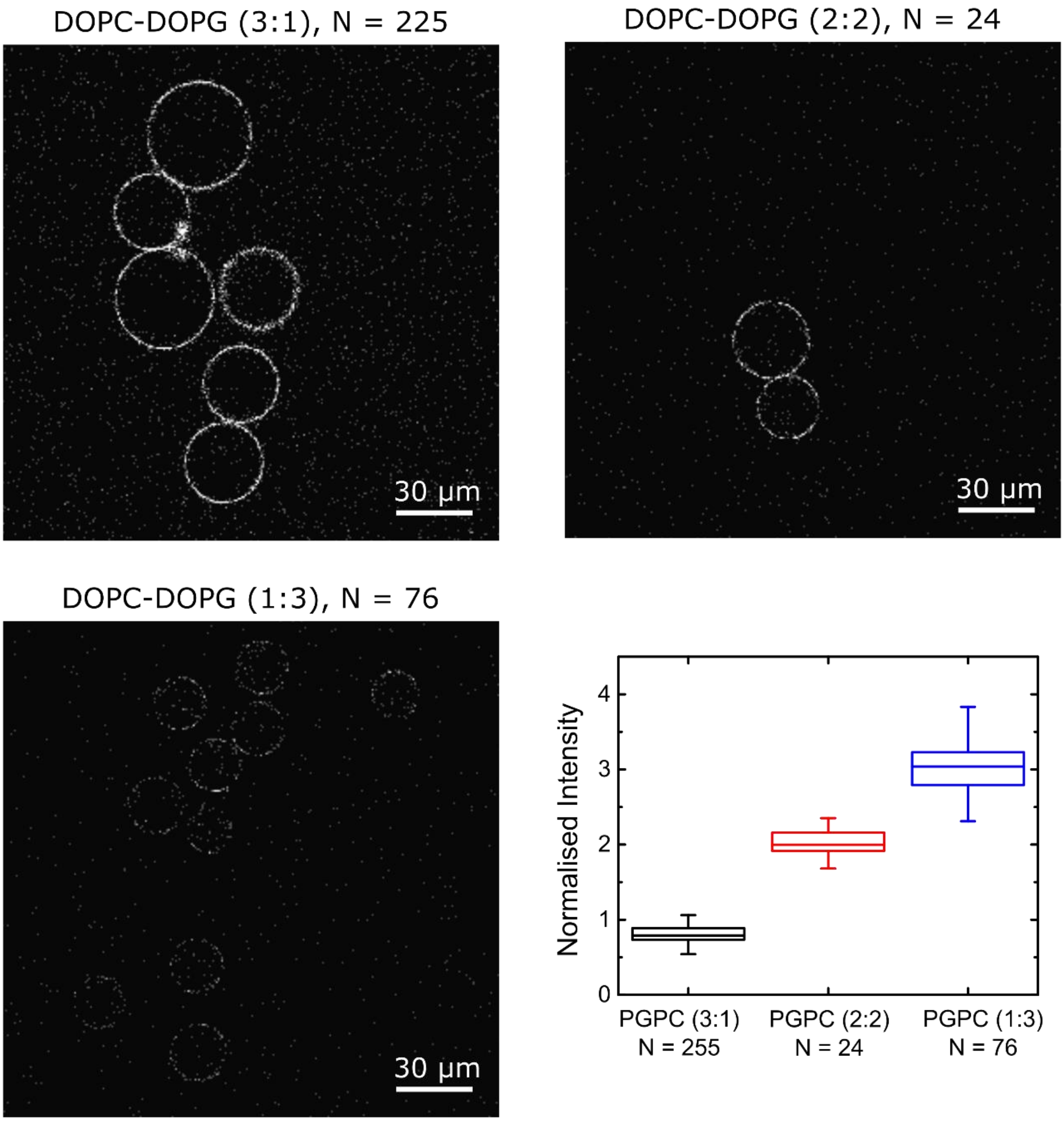


***Figure S2a:*** *Panel showing representative confocal images of PGPC liposomes. The 1:3 ratio vesicles show the highest fluorescence intensity, whereas the 3:1 vesicles show the lowest fluorescence intensity. The vesicles with a 2:2 lipid ratio lie between the two. The boxplot shows the results of the mean fluorescence intensity analysis. The intensity scales in a linear manner in accordance with the relative concentration of fluorescently labelled NBD-PC in the LO phase.*

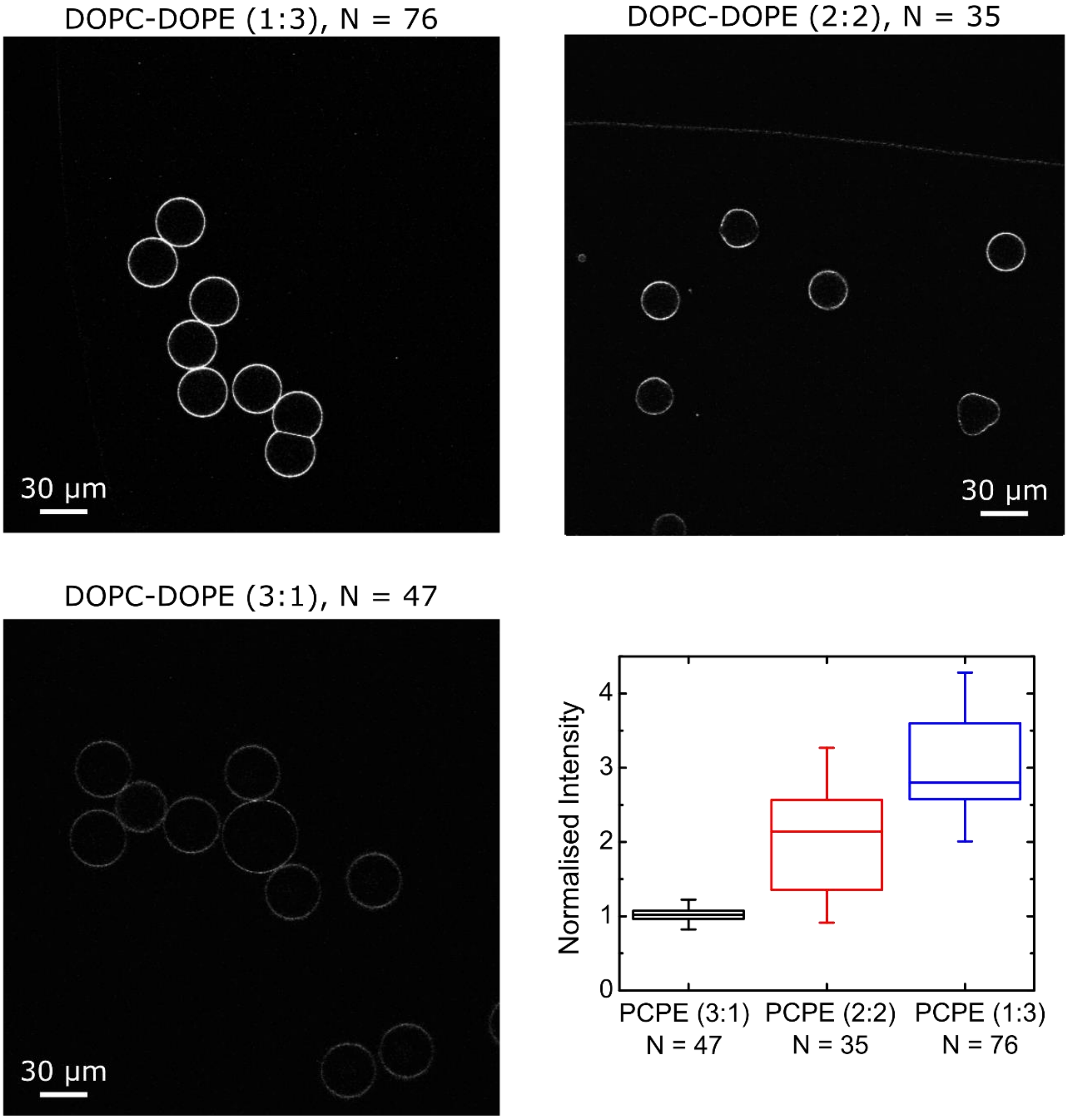


***Figure S2b:*** *Panel showing representative confocal images of PCPE liposomes. The 1:3 ratio vesicles show the highest fluorescence intensity, whereas the 3:1 vesicles show the lowest fluorescence intensity. The vesicles with a 2:2 lipid ratio lie between the two. The boxplot shows the results of the mean fluorescence intensity analysis. The intensity scales in a linear manner in accordance with the relative concentration of fluorescently labelled Liss Rhod PE in the LO phase.*

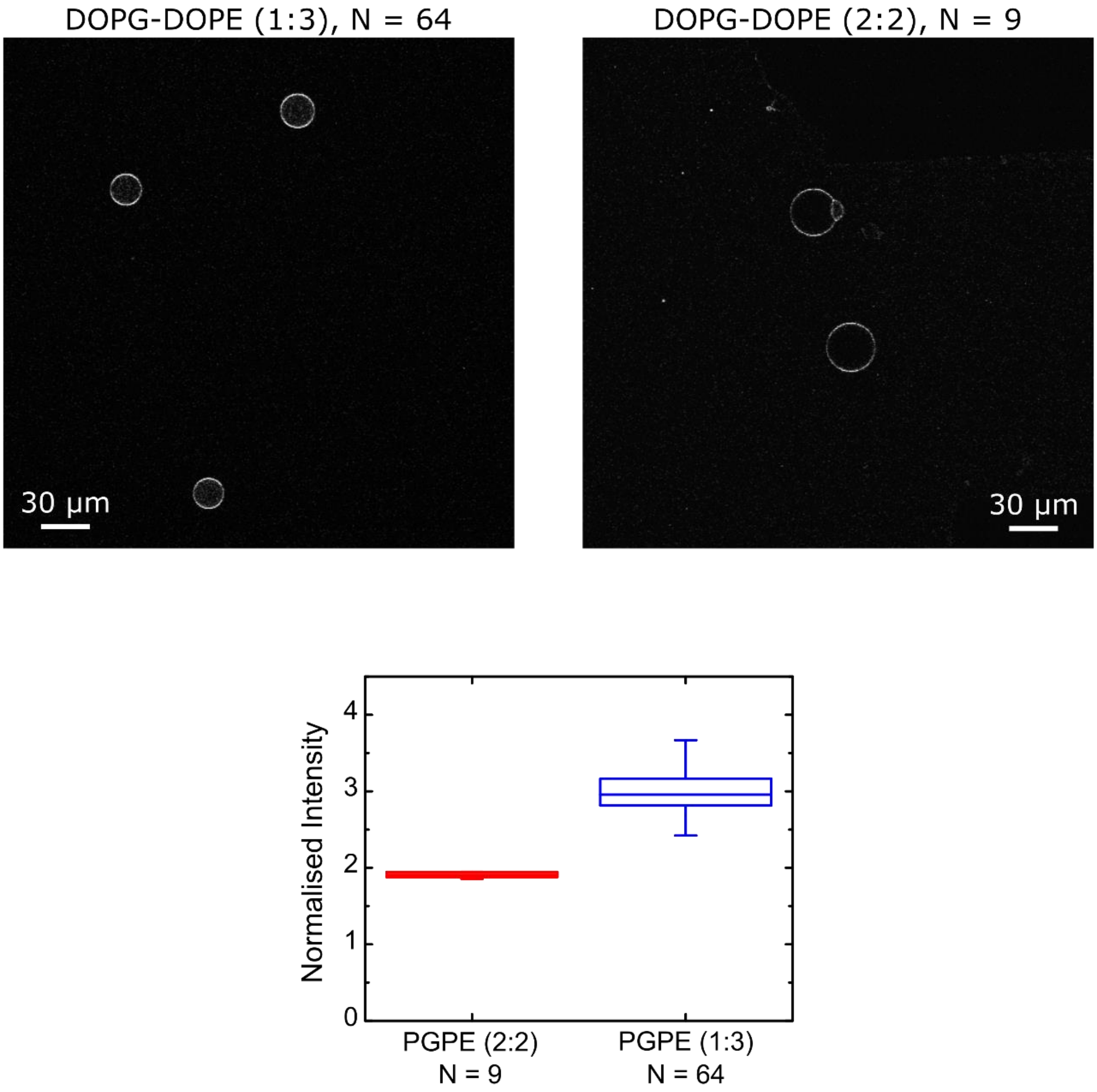


***Figure S2c:*** *Panel showing representative confocal images of PGPE liposomes. The 1:3 ratio vesicles show the highest fluorescence intensity, whereas the 2:2 vesicles show lower fluorescence intensity. We were not able to form stable PGPE vesicles in 3:1 ratio. However, the fluorescence intensity of the other two lipid mixtures scales in a linear manner in accordance with the relative concentration of fluorescently labelled Liss Rhod PE in the LO phase.*

### 2. Lateral Diffusion Experiments

#### 2.1. Comparison of GUV production technique at varying P-188 concentrations

The composition of the solutions used to prepare the vesicles are listed in Table S1. We formed GUVs via electroformation in two chemical environments. One environment was devoid of the poloxamer P-188 in the vesicle solution ('no P-188 environment'), and one environment contained 50 mg/mL P-188 ('high P-188 environment'). Similarly, we formed OLA vesicles in two chemical environments with varying P-188 concentrations. In one case, we prepared GUVs with 50 mg/mL P-188 encapsulated within (IA) and outside (OA) of the vesicles ('high P-188 environment'). However, since OLA vesicles cannot be formed without the presence of P-188 we were not able to create an environment completely devoid of P-188. Instead we formed the OLA GUVs in a 'low P-188 environment' where the inner aqueous (IA) solution contained no P-188 and the outside solution contained 50 mg/mL. The vesicle aliquots had to be further diluted for imaging. The exact solution compositions in which the FRAP measurements were performed are listed in Table S2.

| | Inner Aqueous (IA) | Outer Aqueous (OA) | Dilution stock |
|---|---|---|---|
| **Electroformation** (no P-188 environment) | - 200 mM sucrose<br>- 15% v/v glycerol | - 200 mM sucrose<br>- 15% v/v glycerol | - 200 mM glucose<br>- 15% v/v glycerol |
| **Electroformation** (high P-188 environment) | - 200 mM sucrose<br>- 15% v/v glycerol<br>- 50 mg/mL P-188 | - 200 mM sucrose<br>- 15% v/v glycerol<br>- 50 mg/mL P-188 | - 200 mM glucose<br>- 15% v/v glycerol |
| **OLA** (low P-188 environment) | - 200 mM sucrose<br>- 15% v/v glycerol | - 200 mM sucrose<br>- 15% v/v glycerol<br>- 50 mg/mL P-188 | - 200 mM glucose<br>- 15% v/v glycerol |
| **OLA** (high P-188 environment) | - 200 mM sucrose<br>- 15% v/v glycerol<br>- 50 mg/mL P-188 | - 200 mM sucrose<br>- 15% v/v glycerol<br>- 50 mg/mL P-188 | - 200 mM glucose<br>- 15% v/v glycerol |

***Table S1:*** *Solution compositions used for GUV formation. After production, 20 µL of the vesicle stock solution is extracted and diluted in 50 µL of glucose solution. The lower density of the surrounding medium causes the GUVs to sink to the bottom of the incubation chamber which facilitates confocal imaging. The components were mixed in ion free milli-Q water, as this increases liposome yield during electroformation.*

For visualization, we mixed 20 µL of the vesicle stock with 50 µL of the low-density dilution stock in an incubation chamber (Grace Bio-Labs FlexWell, Sigma-Aldrich) with a wide bore pipette. The density mismatch causes the GUVs to sink to the bottom of the chamber, where they could be imaged. However, by diluting the vesicle stock with P-188 free glucose solution, we reduced the effective P-188 concentration in the outside of the vesicles. The effective solution compositions that the FRAP measurements were carried out in are summarized in Table S2.

| | *Encapsulated Solution* | *Outside Solution* |
|---|---|---|
| **Electroformation**<br>(no P-188 environment) | - 200 mM sucrose<br>- 15% v/v glycerol | - 53 mM sucrose<br>- 143 mM glucose<br>- 15% v/v glycerol<br>- 0 mg/mL P-188 |
| **Electroformation**<br>(high P-188 environment) | - 200 mM sucrose<br>- 15% v/v glycerol<br>- 50 mg/mL P-188 | - 53 mM sucrose<br>- 143 mM glucose<br>- 15% v/v glycerol<br>- 14 mg/mL P-188 |
| **OLA**<br>(low P-188 environment) | - 200 mM sucrose<br>- 15% v/v glycerol | - 53 mM sucrose<br>- 143 mM glucose<br>- 15% v/v glycerol<br>- 14 mg/mL P-188 |
| **OLA**<br>(high P-188 environment) | - 200 mM sucrose<br>- 15% v/v glycerol<br>- 50 mg/mL P-188 | - 53 mM sucrose<br>- 143 mM glucose<br>- 15% v/v glycerol<br>- 14 mg/mL P-188 |

***Table S2:*** *Effective solution composition encapsulated within the vesicles and outside at which the FRAP measurements were performed. We measure the lateral lipid diffusion coefficients in a high and a low P-188 environment for both GUV formation techniques. P-188 could not be completely removed from the outside solution for the OLA sets, as this technique requires a certain amount of P-188 for successful liposome formation.*

| | **DOPC** | **POPC** |
|---|---|---|
| **Electroformation**<br>(no P-188 environment) | 1.0 ± 0.2 $\mu m^2/s$<br>N = 17 | 0.8 ± 0.2 $\mu m^2/s$<br>N = 28 |
| **Electroformation**<br>(high P-188 environment) | 1.2 ± 0.4 $\mu m^2/s$<br>N = 14 | 1.3 ± 0.4 $\mu m^2/s$<br>N = 20 |
| **OLA**<br>(low P-188 environment) | 1.1 ± 0.2 $\mu m^2/s$<br>N = 34 | 1.0 ± 0.3 $\mu m^2/s$<br>N = 49 |
| **OLA**<br>(high P-188 environment) | 1.0 ± 0.3 $\mu m^2/s$<br>N = 30 | 0.9 ± 0.3 $\mu m^2/s$<br>N = 27 |

***Table S3****: Summary of measured lateral lipid diffusion coefficients (mean ± std. dev.) for the GUVs investigated. All measured lateral diffusion coefficients are on the order of 1* $\mu m^2/s$.

### 2.2. Effect of Glycerol and Temperature on Lipid Lateral Diffusion

In order to determine the effect of glycerol and temperature on the lateral lipid diffusion coefficient, we conducted FRAP experiments on electroformed DOPC vesicles with varying glycerol content and at two different temperatures. Sucrose/glucose solutions similar to the 'no P-188 environment' with and without 15% glycerol were prepared. FRAP measurements were conducted at room temperature (approx. 20°C) as well as at 37°C, but with the same optical parameters. Temperature control was performed with the cellVivo incubation system. The obtained values are presented in Table S4.

| | **DOPC (electroformed)** |
|---|---|
| Room temperature (approx. 20°C)<br>15% glycerol | 1.0 ± 0.2 $\mu m^2/s$<br>N = 17 |
| Room temperature (approx. 20°C)<br>0% glycerol | 1.6 ± 0.2 $\mu m^2/s$<br>N = 12 |
| 37°C<br>15% glycerol | 1.9 ± 0.6 $\mu m^2/s$<br>N = 19 |
| 37°C<br>0% glycerol | 2.2 ± 0.5 $\mu m^2/s$<br>N = 7 |

***Table S4:*** *Lateral diffusion coefficients (mean ± std. dev) of electroformed DOPC vesicle membranes at different temperatures and varying glycerol concentrations. The diffusion coefficient increases from 1.0 ± 0.2 $\mu m^2/s$ (mean ± std. dev, N = 17) at 20°C and 15% glycerol to 1.6 ± 0.2 $\mu m^2/s$ (N = 12) without the presence of glycerol. At 37°C, the coefficients rise to 1.9 ± 0.6 $\mu m^2/s$ (N = 19) with 15% glycerol and 2.2 ± 0.5 $\mu m^2/s$ without glycerol.*

### 2.3. Confocal Scans of Vesicles

Figure S3 shows an isometric view as well as a sliced view of three vesicles produced with OLA. Two vesicles have a visible octanol pocket attached to their top side. The octanol manifests itself as a bright spot. The FRAP analysis showed no significant difference in lateral lipid diffusion coefficients of vesicles with or without a visible octanol pocket attached.

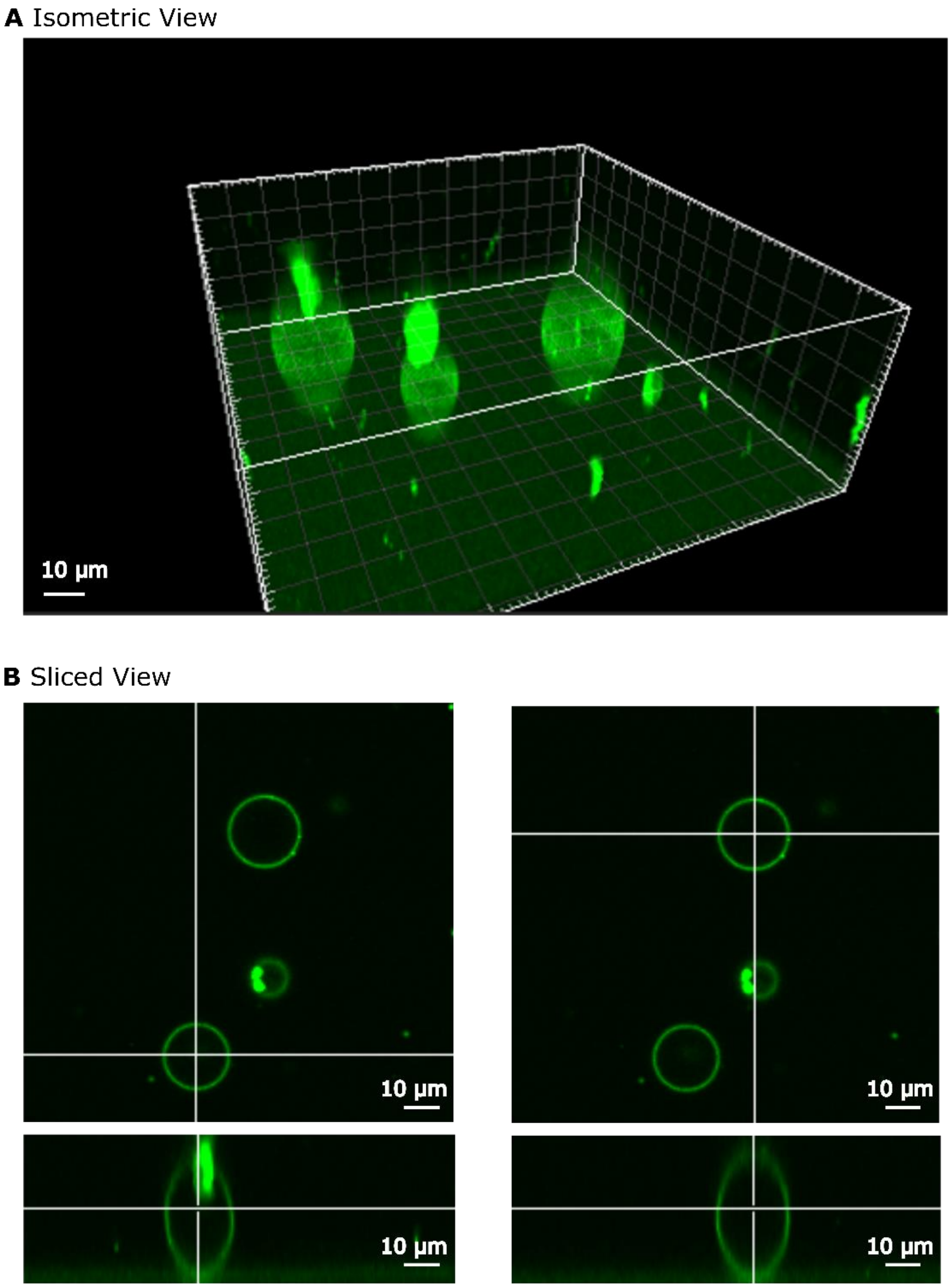


***Figure S3:*** *(A) Isometric view of three GUVs produced with OLA. Two vesicles have a bright octanol pocket attached to their surface. (B) Sliced view of two of the vesicles. The white lines indicate the slice planes. The upper image shows the top view, whereas the bottom image shows the vesicle from the side. A bright octanol pocket can be seen on the side view of the left vesicle. It is not visible in the top view, as it is not within the slice plane. The view on the right shows a vesicle without a visible bright octanol pocket attached to it.*

#### 2.4. Statistical Tests

The following tables show the results of the statistical tests performed on the lateral diffusion coefficients of the vesicles in different chemical environments (Tables S5, S6, S9 and S10), as well as with and without a visible octanol pocket attached (Tables S7, S8, S11 and S12).

##### 2.4.1. DOPC

| | N Analysis | N Missing | Mean | Standard Deviation | SE of Mean |
|---|---|---|---|---|---|
| **Electroformation** (no P-188 environment) | 17 | 0 | 1.0066 | 0.18878 | 0.04579 |
| **Electroformation** (high P-188 environment) | 14 | 0 | 1.2347 | 0.40438 | 0.10807 |
| **OLA** (low P-188 environment) | 34 | 0 | 1.10313 | 0.21012 | 0.03604 |
| **OLA** (high P-188 environment) | 30 | 0 | 0.9801 | 0.25758 | 0.04703 |

| | DF | Sum of Squares | Mean Square | F Value | Prob>F |
|---|---|---|---|---|---|
| **Model** | 3 | 0.72666 | 0.24222 | 3.62715 | 0.01593 |
| **Error** | 91 | 6.077 | 0.06678 | | |
| **Total** | 94 | 6.80366 | | | |

| R-Square | Coeff Var | Root MSE | Data Mean |
|---|---|---|---|
| 0.1068 | 0.24233 | 0.25842 | 1.06639 |

***Table S5:*** *ANOVA on lateral lipid diffusion coefficients of DOPC vesicles in the different chemical environments. At the 0.01 level, the population means are not significantly different.*

| Comparison of lateral diffusion coefficient | p-value (2 sample t-test with Welch's correction) |
|---|---|
| Electroformation *'no P-188'* vs. Electroformation *'high P-188'* | 0.06812 |
| Electroformation *'no P-188'* vs. OLA *'low P-188'* | 0.10644 |
| Electroformation *'no P-188'* vs. OLA *'high P-188'* | 0.68838 |
| Electroformation *'high P-188'* vs. OLA *'high P-188'* | 0.04441 |
| Electroformation *'high P-188'* vs. OLA *'low P-188'* | 0.26509 |
| OLA *'low P-188'* vs. OLA *'high P-188'* | 0.04244 |

***Table S6:*** *Two sample t-tests comparing lateral lipid diffusion coefficients of DOPC vesicles in 'no/low' and 'high' P-188 environments. At the 0.01 level, the means are not significantly different from one another.*

| | N Analysis | N Missing | Mean | Standard Deviation | SE of Mean |
|---|---|---|---|---|---|
| **Octanol pocket attached** (high P-188 environment) | 13 | 1 | 0.87757 | 0.17093 | 0.04741 |
| **Octanol pocket separated** (high P-188 environment) | 17 | 2 | 1.0585 | 0.28856 | 0.06999 |

| | DF | Sum of Squares | Mean Square | F Value | Prob>F |
|---|---|---|---|---|---|
| **Model** | 1 | 0.24115 | 0.24115 | 4.01219 | 0.05494 |
| **Error** | 28 | 1.68289 | 0.0601 | | |
| **Total** | 29 | 1.92404 | | | |

| R-Square | Coeff Var | Root MSE | Data Mean |
|---|---|---|---|
| 0.12533 | 0.25014 | 0.24516 | 0.9801 |

***Table S7a:*** *ANOVA on lateral lipid diffusion coefficients of DOPC vesicles in a 'high' P-188 environment with and without a visible octanol pocket attached. At the 0.05 level, the population means are not significantly different.*

| | N Analysis | N Missing | Mean | Standard Deviation | SE of Mean |
|---|---|---|---|---|---|
| **Octanol pocket attached** (low P-188 environment) | 15 | 3 | 1.06813 | 0.20468 | 0.05285 |
| **Octanol pocket separated** (low P-188 environment) | 19 | 2 | 1.13075 | 0.21572 | 0.04949 |

| | DF | Sum of Squares | Mean Square | F Value | Prob>F |
|---|---|---|---|---|---|
| **Model** | 1 | 0.03287 | 0.03287 | 0.73853 | 0.39652 |
| **Error** | 32 | 1.42411 | 0.0445 | | |
| **Total** | 33 | 1.45697 | | | |

| R-Square | Coeff Var | Root MSE | Data Mean |
|---|---|---|---|
| 0.02256 | 0.19124 | 0.21096 | 1.10313 |

***Table S7b:*** *ANOVA on lateral lipid diffusion coefficients of DOPC vesicles in a 'low' P-188 environment with and without a visible octanol pocket attached. At the 0.05 level, the population means are not significantly different.*

| Comparison of lateral diffusion coefficient | **p-value** (2 sample t-test with Welch's correction) |
|---|---|
| OLA octanol pocket attached vs. OLA octanol pocket separated (*'low P-188'*) | 0.39379 |
| OLA octanol pocket attached vs. OLA octanol pocket separated (*'high P-188'*) | 0.04166 |

***Table S8:*** *Two sample t-test comparing lateral lipid diffusion coefficients of DOPC vesicles with and without a visible octanol pocket attached. At 0.01 level, the means are not significantly different from one another.*

#### 2.4.2. POPC

| | N Analysis | N Missing | Mean | Standard Deviation | SE of Mean |
|---|---|---|---|---|---|
| **Electroformation** (no P-188 environment) | 28 | 0 | 0.79702 | 0.20225 | 0.03822 |
| **Electroformation** (high P-188 environment) | 20 | 0 | 1.34993 | 0.43119 | 0.09642 |
| **OLA** (low P-188 environment) | 49 | 0 | 0.97925 | 0.2719 | 0.03884 |
| **OLA** (high P-188 environment) | 27 | 0 | 0.93057 | 0.13276 | 0.02555 |

| | DF | Sum of Squares | Mean Square | F Value | Prob>F |
|---|---|---|---|---|---|
| **Model** | 3 | 3.73392 | 1.24464 | 1.73E+01 | 2.17E-09 |
| **Error** | 120 | 8.64406 | 0.07203 | | |
| **Total** | 123 | 12.37798 | | | |

| R-Square | Coeff Var | Root MSE | Data Mean |
|---|---|---|---|
| 0.30166 | 0.27185 | 0.26839 | 0.98729 |

***Table S9:*** *ANOVA on lateral lipid diffusion coefficients of POPC vesicles in the different chemical environments. At the 0.001 level, the difference in the population means are statistically significant.*

| Comparison of lateral diffusion coefficient | p-value (2 sample t-test with Welch's correction) |
|---|---|
| Electroformation *'no P-188'* vs. Electroformation *'high P-188'* | 1.58549E-5 |
| Electroformation *'no P-188'* vs. OLA *'low P-188'* | 0.00133 |
| Electroformation *'no P-188'* vs. OLA *'high P-188'* | 0.00559 |
| Electroformation *'high P-188'* vs. OLA *'high P-188'* | 3.76001E-4 |
| Electroformation *'high P-188'* vs. OLA *'low P-188'* | 0.00147 |
| OLA *'low P-188'* vs. OLA *'high P-188'* | 0.29855 |

***Table S10:*** *Two sample t-tests comparing lateral lipid diffusion coefficients of POPC vesicles in 'no/low' and 'high' P-188 environments. At the 0.001 level, the lateral lipid diffusion coefficient of electroformed vesicles in a 'high P-188' environment is significantly different from electroformed vesicles without P-188, as well as OLA vesicles with 'high P-188'. All remaining diffusion coefficients are not significantly different from one another.*

| | N Analysis | N Missing | Mean | Standard Deviation | SE of Mean |
|---|---|---|---|---|---|
| **Octanol pocket attached** (high P-188 environment) | 17 | 1 | 0.94456 | 0.13026 | 0.03159 |
| **Octanol pocket separated** (high P-188 environment) | 9 | 2 | 0.88763 | 0.13453 | 0.04484 |

| | DF | Sum of Squares | Mean Square | F Value | Prob>F |
|---|---|---|---|---|---|
| **Model** | 1 | 0.01907 | 0.01907 | 1.09968 | 0.30478 |
| **Error** | 24 | 0.41627 | 0.01734 | | |
| **Total** | 25 | 0.43534 | | | |

| R-Square | Coeff Var | Root MSE | Data Mean |
|---|---|---|---|
| 0.04381 | 0.1424 | 0.1317 | 0.92485 |

***Table S11a:*** *ANOVA on lateral lipid diffusion coefficients of POPC vesicles in a high P-188 environment with and without a visible octanol pocket attached. At the 0.05 level, the population means are not significantly different.*

| | N Analysis | N Missing | Mean | Standard Deviation | SE of Mean |
|---|---|---|---|---|---|
| **Octanol pocket attached** (low P-188 environment) | 32 | 11 | 0.93381 | 0.28302 | 0.05003 |
| **Octanol pocket separated** (low P-188 environment) | 17 | 5 | 1.06477 | 0.23388 | 0.05672 |

| | DF | Sum of Squares | Mean Square | F Value | Prob>F |
|---|---|---|---|---|---|
| **Model** | 1 | 0.19039 | 0.19039 | 2.66451 | 0.10929 |
| **Error** | 47 | 3.35829 | 0.07145 | | |
| **Total** | 48 | 3.54868 | | | |

| R-Square | Coeff Var | Root MSE | Data Mean |
|---|---|---|---|
| 0.05365 | 0.27297 | 0.26731 | 0.97925 |

***Table S11b:*** *ANOVA on lateral lipid diffusion coefficients of POPC vesicles in a low P-188 environment with and without a visible octanol pocket attached. At the 0.05 level, the population means are not significantly different.*

| Comparison of lateral diffusion coefficient | p-value (2 sample t-test with Welch's correction) |
|---|---|
| OLA octanol pocket attached vs. OLA octanol pocket separated (*'low P-188'*) | 0.09139 |
| OLA octanol pocket attached vs. OLA octanol pocket separated (*'high P-188'*) | 0.25367 |

***Table S12:*** *Two sample t-tests comparing lateral lipid diffusion coefficients of POPC vesicles with and without a visible octanol pocket attached. At 0.05 level, the means are not significantly different from one another.*
